# Cuticular Wax Composition is Essential for Plant Recovery Following Drought with Little Effect under Optimal Conditions

**DOI:** 10.1101/2021.06.08.447487

**Authors:** Boaz Negin, Shelly Hen-Avivi, Efrat Almekias-Siegl, Lior Shachar, Asaph Aharoni

**Affiliations:** Department of Plant and Environmental Sciences, Weizmann Institute of Science, Rehovot, Israel

## Abstract

Despite decades of extensive study, the role of cuticular lipids in sustaining plant fitness is far from being understood. To answer this fundamental question, we employed genome editing in tree tobacco (*Nicotiana glauca*) plants and generated mutations in 16 different cuticular lipids-related genes. We chose tree tobacco due to the abundant, yet simply composed epicuticular waxes deposited on its surface. Five out of 9 different mutants that displayed a cuticular lipids-related phenotype were selected for in depth analysis. They had either reduced total wax load or complete deficiency in certain wax components. This led to substantial modification in surface wax crystal structure and to elevated cuticular water loss. Remarkably, under non-stressed conditions, mutant plants with altered wax composition did not display elevated transpiration or reduced growth. However, once exposed to drought, plants lacking alkanes were not able to strongly reduce their transpiration, leading to leaf death and impaired recovery upon resuscitation, and even to stem cracking, a phenomenon typically found in trees experiencing drought stress. In contrast, plants deficient in fatty alcohols exhibited an opposite response, having reduced cuticular water loss and rapid recovery following drought. This differential response was part of a larger trend, of no common phenotype connecting plants with a glossy appearance. We conclude that alkanes are essential under drought response and much less under normal non-stressed conditions, enabling plants to seal their cuticle upon stomatal closure, reducing leaf death and facilitating a speedy recovery.

## Introduction

Plants aerial organs are exposed to a plethora of conditions that require a barrier to shield their inner tissues. Such layers protect them from many factors including a desiccating environment, high radiation in different wavelengths and colonization or feeding by other organisms such as fungi, bacteria, insects and even grazing mammals. To cope with these, plants developed a complex surface layer, which at times may also contain specialized components that aid the plant in its response to the different factors. The cuticle may be imbedded with flavonoids (Ryan et al., 2001; Ryan et al., 2002) and anthocyanins (Chalker-Scott, 1999) reducing UVB damage to the mesophyll. Trichomes present on the surface may serve as a structural and chemical obstacle for plant herbivores (Hanley et al., 2007) and change leaf reflectance, reducing photoinhibition and UV-B related damage (Steffens and Walters, 1991; Peiffer et al., 2009; Sonawane et al., 2020; Bickford, 2016). The plant cuticle consists of three core elements localized beyond the epidermis cells. The cutin polymer composed of an amorphous matrix largely comprising C16 and C18 fatty acids connected in esteric bonds (Yeats and Rose, 2013), intracuticular wax embedded in the cutin layer and epicuticular waxes secreted beyond the cutin. Though it is clear that the cuticle plays a crucial role in preventing non stomatal water loss, the extent to which epicuticular wax contributes to this varies widely and in some cases may be negligible compared to the other two elements (Jetter and Riederer, 2016; Zeisler and Schreiber, 2016; Zeisler-Diehl et al., 2018). This keeps open the question of epicuticular wax’s function, in cases where its contribution to the transpirational barrier in small. A related question, is to what extent the different wax components contribute to the different functions of epicuticular waxes.

Decades of extensive research, mainly in the model species *Arabidopsis thaliana* resulted in elucidation of most biosynthetic pathways associated with production of the different wax components (Fig. 1; Lee and Suh, 2015). In Arabidopsis, fatty acids synthesized in plastids are transported to the endoplasmic reticulum (ER) where they are converted to acyl CoA by LONG CHAIN ACYL SYNTHETASE (LACS; Schnurr et al., 2004; Lü et al., 2009). The fatty acid elongase complex composed of four proteins then elongates these C16-C18 precursors. These include (i) β-ketoacyl-CoA synthase (KCS; Millar and Kunst, 1997) having different variants responsible for elongation of certain acyl substrates (e.g. KCS6 targeted here is responsible for elongation from 24 to 34 carbons (Millar et al., 1999), (ii) β-ketoacyl-CoA reductase (KCR; Beaudoin et al., 2009), (iii) 3-hydroxyacyl-CoA dehydratase (HCD; Bach et al., 2008) and (iv) trans-2,3-enoyl-CoA reductase (ECR; Zheng et al., 2005). These enzymes elongate the Acyl CoAs through a cycle in which malonyl CoA and the acyl are condensed and at its end the acyl is elongated by two carbons. The outcome of the elongation process are very long chain fatty (VLCF) acyl CoA of different lengths used in several epicuticular wax biosynthesis pathways. They can be converted into very long chain fatty acids (VLCFA) or enter two paths, the alkane- and alcohol-forming pathways. In the alkane-forming pathway, VLCF acyl-CoAs are decarboxylated to create aldehydes and then alkanes. The exact enzymatic mechanisms taking place during the conversion of acyl-CoA to alkanes are not clear, though it is known that ECERIFERUM1 (CER1) and ECERIFERUM 3 (CER3) take part in the first stages of conversion to alkanes (Aarts et al., 1995; Chen et al., 2003; Bourdenx et al., 2011). Following the synthesis of alkanes, these can be further modified to form secondary alcohols and ketones, by the MID-CHAIN ALKANE HYDROXYLASE1 (MAH1); a cytochrome p450 catalyzing both reactions (Greer et al., 2007). In the primary alcohol forming pathway, VLCF acyl-CoAs are converted to primary alcohols through the addition of a hydroxyl at the acyls end. This step is catalyzed by the FATTY ACYL-COA REDUCTASE (FAR) enzyme (Rowland et al., 2006). An additional stage in the alcohol forming pathway is the conjunction of C16 or C18 fatty acids to the primary alcohols to create wax esters by the bifunctional wax synthase/ acyl CoA:diacylglycerol acyltransferase1 (WSD1; Li et al., 2008). Although the presence of epicuticular wax is almost ubiquitous among plants, there is great diversity between species and even different organs of the same plant, in chemical structure and form of wax crystals they produce (Barthlott et al., 1998; Lee and Suh, 2015).

**Figure 1.**
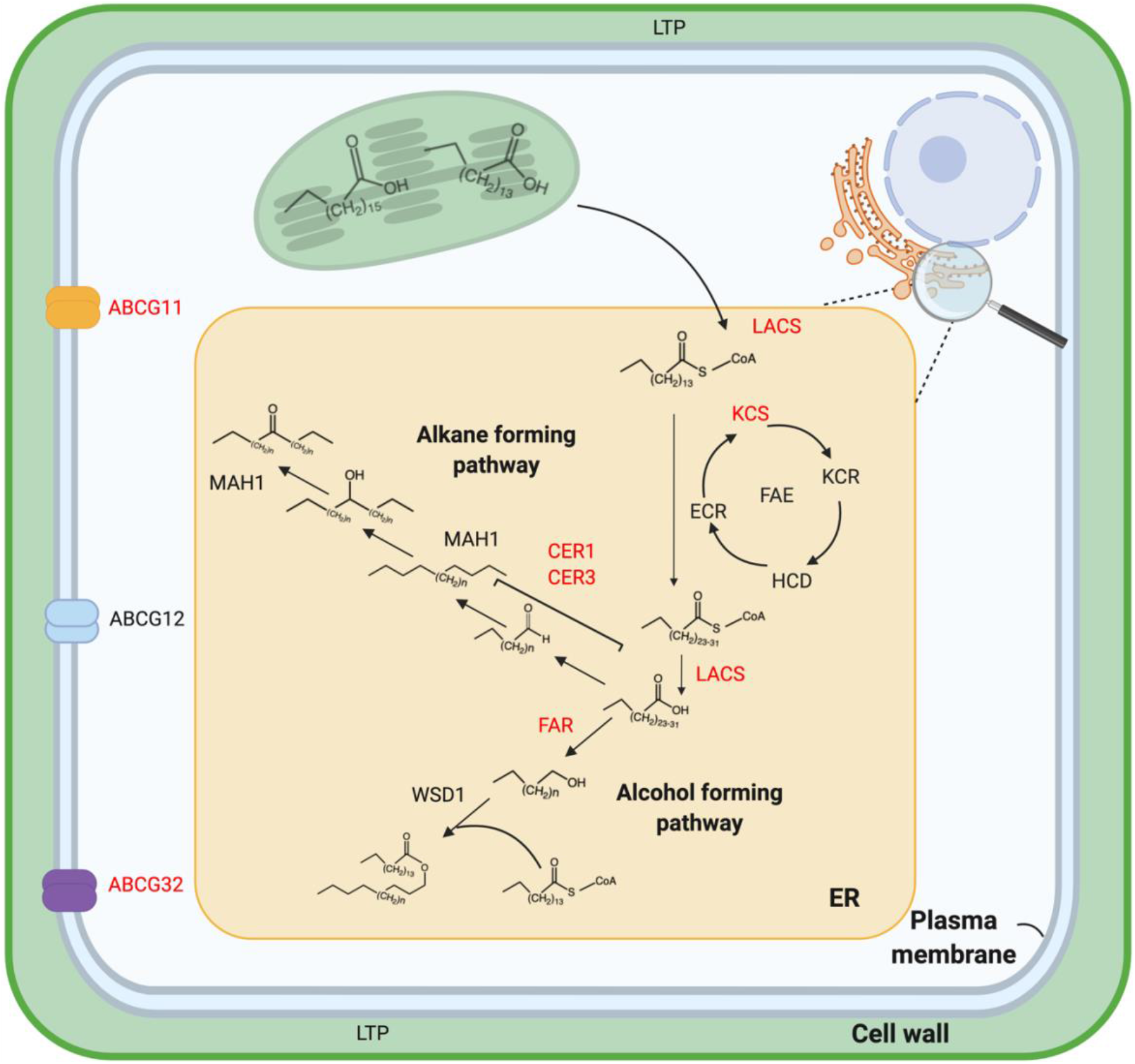
A schematic model of epicuticular wax biosynthesis and transport pathways. Not all proteins involved in these pathways are presented and genes corresponding to proteins marked in red were targeted for editing and exhibited a visual phenotype in this study. The scheme presents fatty acid elongation and division to the two main wax synthesis pathways – the alkane and fatty alcohol forming pathways. Genes silenced that are involved in wax synthesis but not a specific pathway include *KCS6* and *LACS.* Those taking part in alkane synthesis are *CER1*, *CER3* and *MAH1*, and fatty alcohol synthetic genes include *FAR* and *WSD1*.

Tree tobacco (*Nicotiana glauca*) is a perineal shrub originating in South America which has since spread worldwide. The appearance of its stems and leaves is glaucous, an uncommon appearance in tobacco species which gave the species its name. This glaucous appearance, is the result of a high load of epicuticular wax coating the plant’s aerial organs. This wax is composed almost solely of C_31_ alkanes with lesser amounts of additional alkanes, fatty alcohols and aldehydes (Mortimer et al., 2012). Furthermore, *N. glauca* accumulates substantial amounts of wax in response to drought, while maintaining its original composition (Cameron et al., 2006). These characteristics including the availability of its genome sequence (Usadel et al., 2018), efficient stable transformation and the ease of dissecting its epidermis layer make it an excellent model plant for studying the yet unresolved role of epicuticular waxes in the plant life cycle.

In this study, we employed CRISPR-Cas9 based technology to edit 16 cuticular lipids metabolism genes and generate mutants in *N. glauca*. Following initial characterization, we carried out in-depth research on five selected knockout mutants that showed diverse patterns of wax composition. This mutant set was subjected to a range of examinations to help us better understand what the contribution of epicuticular wax is to plant fitness, and to attribute this contribution to specific wax components. We found, that under optimal conditions epicuticular wax has little effect on plant fitness. In contrast, following drought the alkane fraction is essential for plant recovery, whereas deficiency in fatty alcohols did not affect plants negatively under our experimental conditions. Findings in this study highlight the specific role of plant epicuticular wax in episodes of drought and the following recovery phase. They further explain how diversity of wax components and structures contribute to plant fitness under abiotic stress conditions.

## Results

### Transcriptomics in *Nicotiana glauca* epidermis tissues under drought conditions facilitates the discovery of cuticular lipids-related genes

The most striking feature of *N. glauca* is the glaucous waxy appearance formed by an extremely high but very simply composed wax layer covering its above ground organs. Such intense wax coverage provokes a fundamental question with respect to the high carbon investment in epicuticular wax production set against its contribution to plant fitness. Which wax component contributes to fitness and in what manner is a consequent key question. To answer these, it was first essential to identify cuticular lipids-associated genes active in the *N. glauca* epidermis layer. We thus performed transcriptomic analysis of five different shoot tissues including dissected adaxial and abaxial leaf epidermis, stem epidermis, complete leaves, and whole stems. Expression profiling was carried out in well-watered plants or those under drought conditions, as a previous study demonstrated drought-induced wax formation in *N. glauca* leaves (Cameron et al., 2006). To detect genes associated with cuticular lipids formation we next mined the transcriptome dataset for transcripts that were epidermis enriched, drought induced or a combination of the two conditions. We found that many homologs of cuticular lipids metabolism genes (e.g. *KCS6*, *CER1*, *FAR* and *ABCG32*), displayed a similar ‘expression signature’; high epidermal enrichment and mild drought induction (Fig. 2). Out of tens of genes putatively associated with cuticular lipids metabolism we selected 16 for further study. Editing these genes could alter wax and cutin synthesis and transport in a variety of manners resulting in a set of mutant lines with diverse was compositions (Table S1). This set comprised seven genes putatively encoding wax biosynthesis enzymes including *LACS1*, *KCS6*, *CER1*, *CER3*, *MAH1*, *FAR* and *WSD1* and three cutin biosynthesis enzymes - Glycerol-3-phosphate 2-O-acyltransferase 4 (*GPAT4*), Cytochrome P450 86A22 (*CYP86A22*) and GDSL-motif esterase/acyltransferase/lipase (*GDSL*). Three putative transporters - ATP-BINDING CASSETTE G11 (*ABCG11*), ATP-BINDING CASSETTE G32 (*ABCG32*) and ACYL-COA-BINDING PROTEIN 1 (*ACBP1*) and three transcription factor homologs - *SHINE1*, *SHINE3* and *MYB96*, complemented the set of selected genes. We next generated plant transformation vectors for editing the 16-gene set through CRISPR-Cas9 technology and introduced them to *N. Glauca* (Fig. S2-S10).

**Figure 2.**
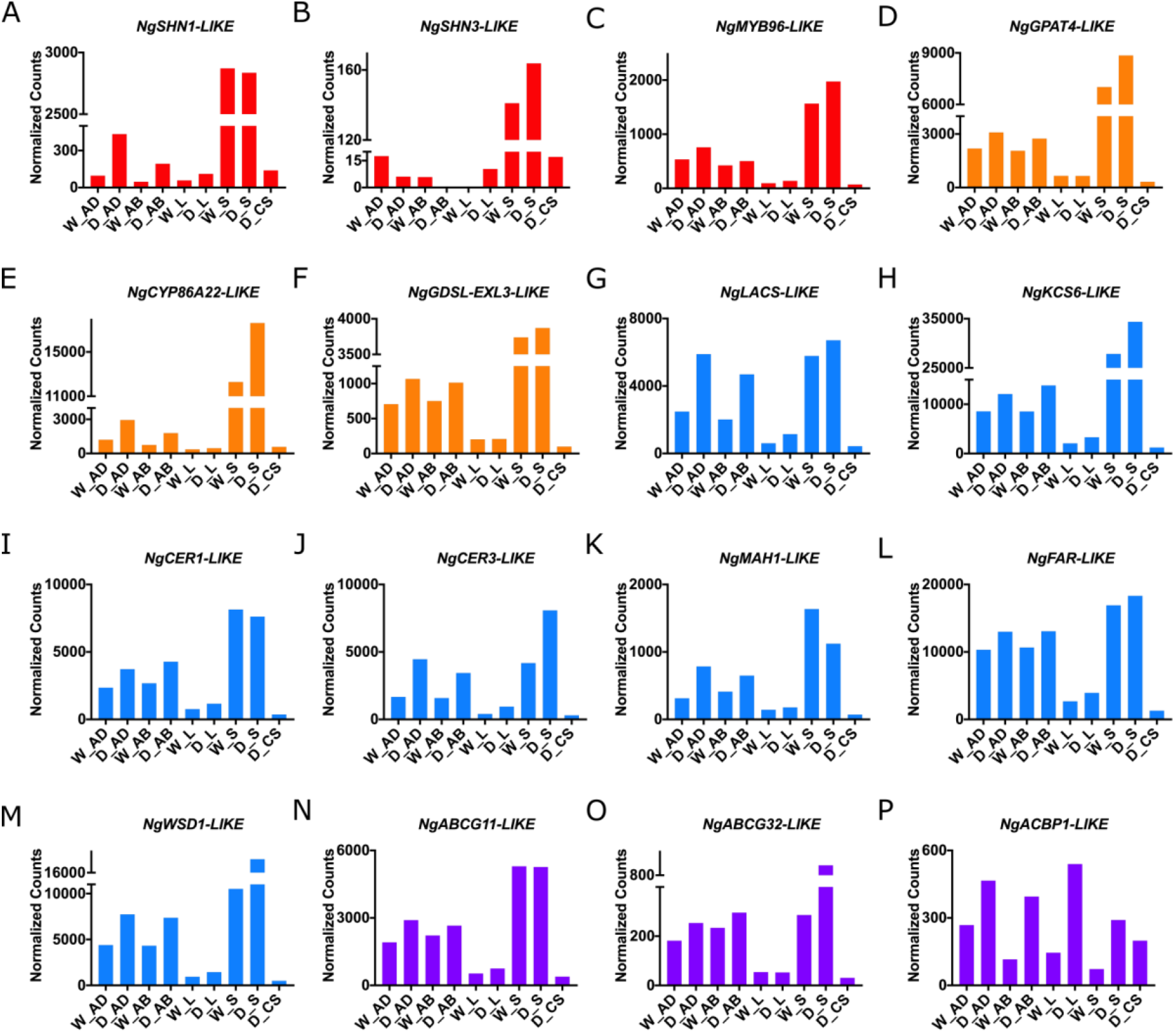
Epidermis enriched and drought-induced expression of *N. Glauca* genes showing homology to known cuticular lipids genes from other species. For each sample, three biological replicates of each tissue, grown either under well-watered conditions or drought treated, were collected and pooled to reduce variance though not increasing the number of sequenced samples. The different genes are color coded according to their function; transcription factors in red **(A-C),** cutin synthesis genes in orange **(D-F),** wax synthesis genes in blue **(G-M),** and transporter genes in purple **(N-P)**. AD- adaxial epidermis tissue; AB- abaxial epidermis tissue; L- whole leaf tissue; S- stem epidermis tissue; CS-complete stem; W- watered plants; D- drought treated plants.

### Cuticular lipids mutants display diverse surface and morphological phenotypes

Of the 16 genes targeted, nine exhibited visual surface related phenotypes at the T0 generation (Fig. 3 and Fig. S1). These included glossy leaves and stems and fused anthers in the *kcs6* mutant alleles, glossy leaves and stems in the *far* and *cer1* mutants and glossy leaves alongside waxy stems in *cer3* mutants (Fig. 3). The *abcg11* mutant, the only line that never flowered (even over two years) possessed small, glossy, crinkled, brittle leaves. Both *cyp86a22* and *abcg32* showed elongated and malformed leaves, while *gpat4* displayed elongated leaves and leaf fusions. Finally, the *lacs* mutants displayed both a glossy phenotype and leaf deformities (Fig. S1). Sequencing the amplicons covering the targeted editing sites revealed a wide range of mutations; from single base pair insertions or deletions, deletion of a triplet coding for a single amino acid to a 200 bp deletion (Fig S2-S10). First generation plants (i.e. T0) had both homozygous and heterozygous mutations, and frequently carried two different mutations on their two chromosomes, and not merely two identical mutations.

**Figure 3.**
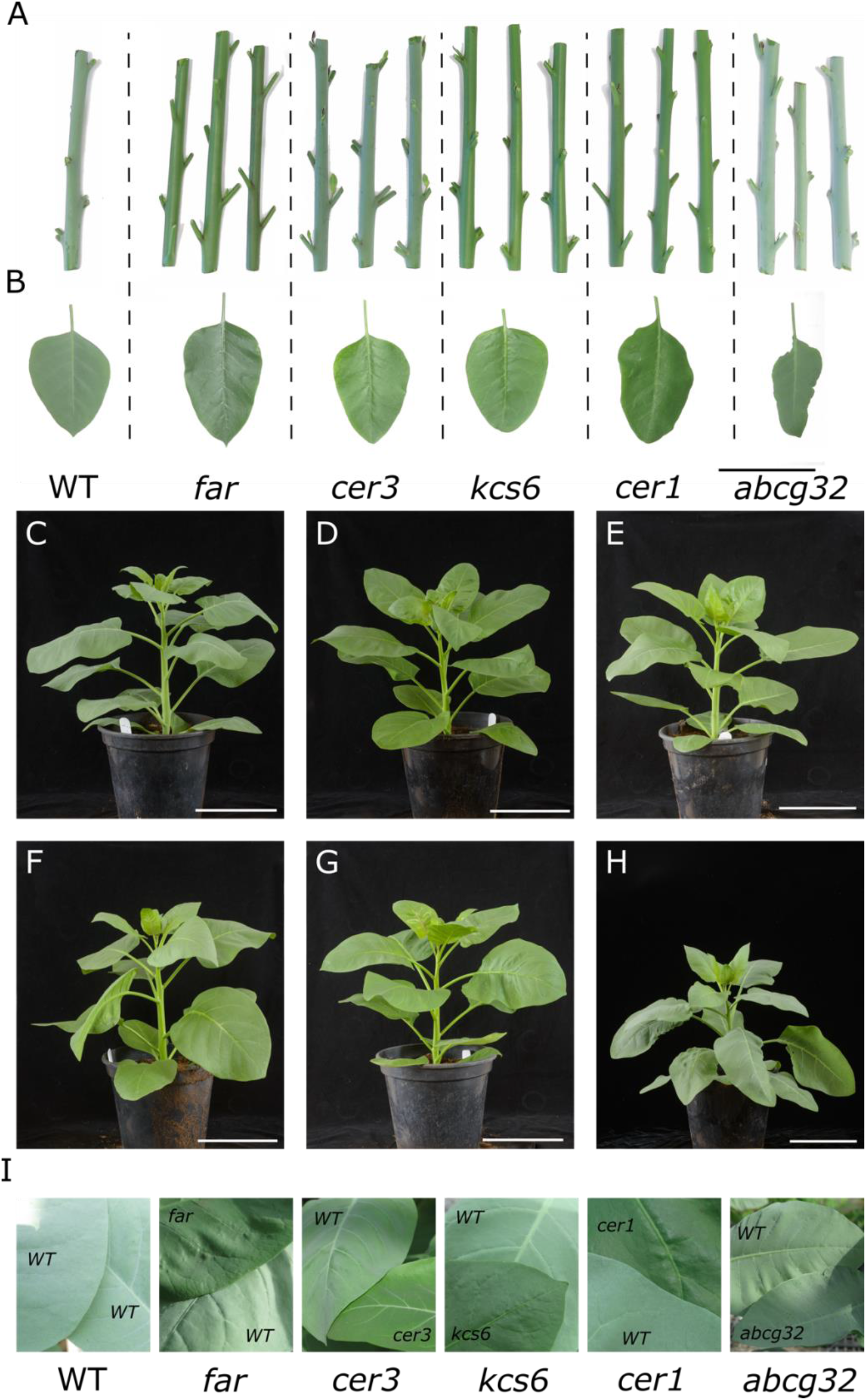
Plant surface phenotypes of mutant plants analyzed in this study. **(A)** Representative stems of wild type (WT) and the five independent mutant lines investigated in-depth. **(B)** Representative leaves of WT and mutants’ leaves. Bar=5cm. **(C-H)** representative whole plants Bar=15cm. C, WT; D, *far*; E, *cer3*; F, *kcs6*; G, *cer1*; H, *abcg32*. **(I)** images of WT leaves and those of mutants alongside.

### Surface wax composition and crystal morphology in wax genes mutants

Out of the nine mutants, we focused the study on four wax metabolism genes, as well as the *abcg32* mutant that served as a control as it is affected in cutin monomer transport. We next extracted leaves’ epicuticular wax and analyzed its composition using gas chromatography - mass spectrometry (GC-MS). Wax composition in *N. Glauca* is typically dominated by a C_31_ alkane (92% according to Mortimer et al., 2012) along with C33 alkanes, C_24_, C_26_ and C_28_ fatty alcohols and smaller amounts of a C_26_ aldehydes (as well as trace amounts of other alkanes, alcohols and aldehydes). Mutations in the different genes had a drastic effect on epicuticular wax composition (Fig. 4). *kcs6* mutant leaves displayed almost complete reduction in all wax components with a chain length above 26 carbons making them nearly free of alkanes (Fig. 4A-B). The same leaves accumulated shorter chain length waxes such as C18, C20, C22 and C_24_ alcohols (Fig. 4C-F). Leaves of the *cer1* mutants showed massive reduction in abundance of all alkanes (Fig 4A-B) mirrored by a significant increase in C_26_ alcohols (Fig. 4E). *cer3* mutants had a significant reduction (approximately 80%) in their C_31_ wax load. In the same mutant leaves, the C33 alkane showed a trend of increase that was significant in one independent line (Fig. 4A-B). The *far* mutants displayed extremely reduced fatty alcohol content (Fig. 4C-F) contrasted by a trend of increase in their alkane load (Fig 4A-B). *abcg32* mutants’ wax composition was similar to WT, with only a trend of reduction in C_31_ alkane load (Fig. 4A) and a significant increase in C_26_ alcohols in *abcg32-18* and an increase in C_28_ alcohols in both *abcg32-6* and *abcg32-18* (Fig. 4E-F).

**Figure 4.**
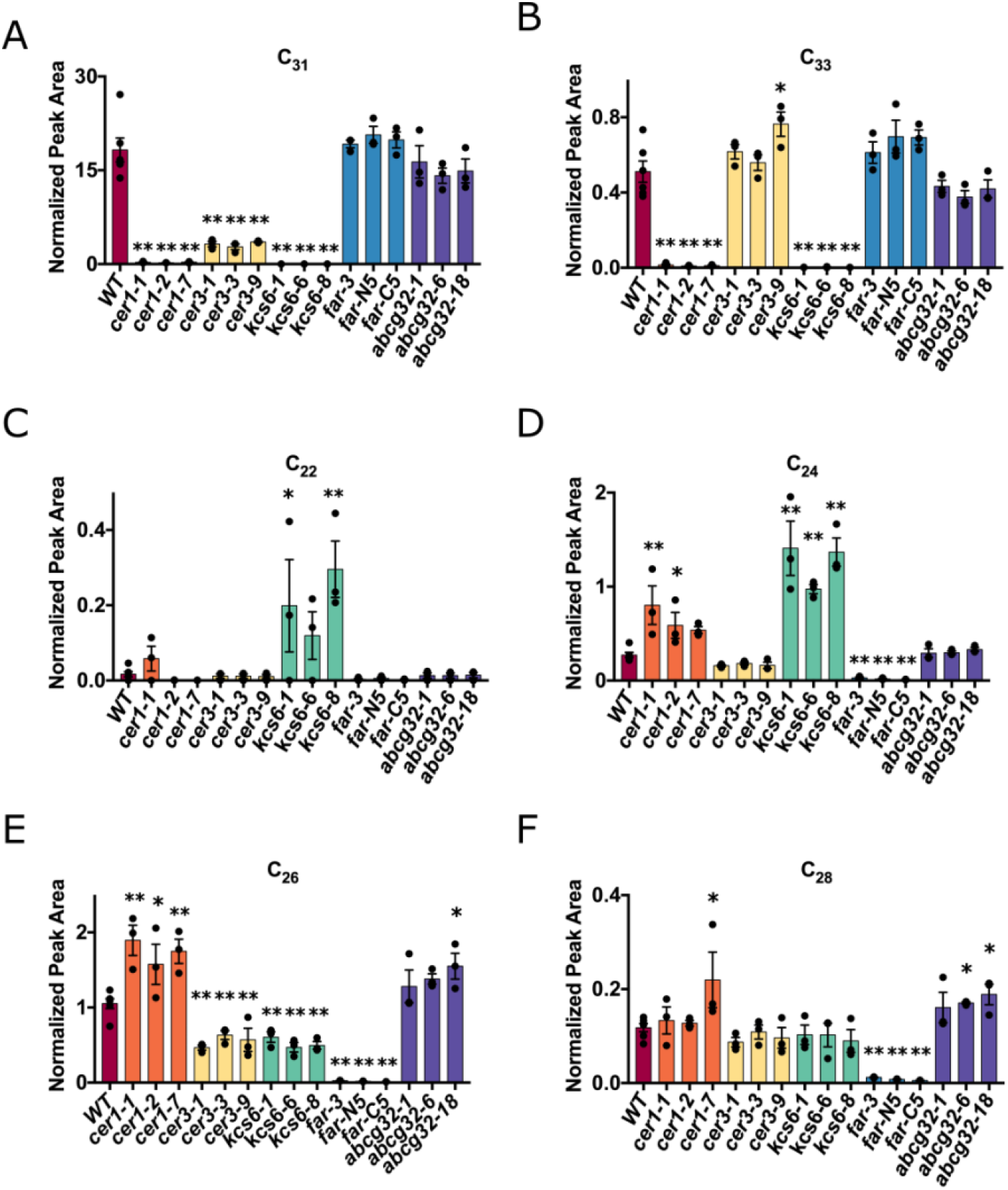
Relative abundance of major wax components in *N. glauca* leaves. **(A-B)** Alkanes and **(C-F)** primary alcohols. Numbers indicate carbon chain length. Independent mutant alleles of the same gene are indicated by the same color. WT, n=6; all other lines n=3. *= p<0.05 and **= p<0.01 as determined in a student’s t test. Bars represent standard error. Wax components were extracted in chloroform, derivatized and quantified by GC-MS

Alterations in composition of epicuticular waxes had a strong effect on wax crystal morphology as visualized using cryo-SEM (Fig. 5). The typical wax crystals of *N. Glauca* leaves are of dense rodlets (Fig. 5A). In *cer1* leaves, a thin layer of flakes replaced wax rodlets normally found in WT leaves (Fig. 5B). Leaves of *cer3* displayed crystal morphology similar to that of the WT leaves, however, these crystals were reduced in size and more sparsely distributed on the leaf surface (Fig. 5C). Leaf wax of *kcs6* lost its rodlet-like morphology and instead appeared as structured lines of vertically positioned membranous platelets (Fig. 5D). The *far* leaves exhibited large vertical sharp-edged plates, which were sparsely distributed on the leaf (Fig. 5E). Finally, the *abcg32* mutant wax crystals appeared normal as expected from its almost unaltered wax composition (Fig. 5F).

**Figure 5.**
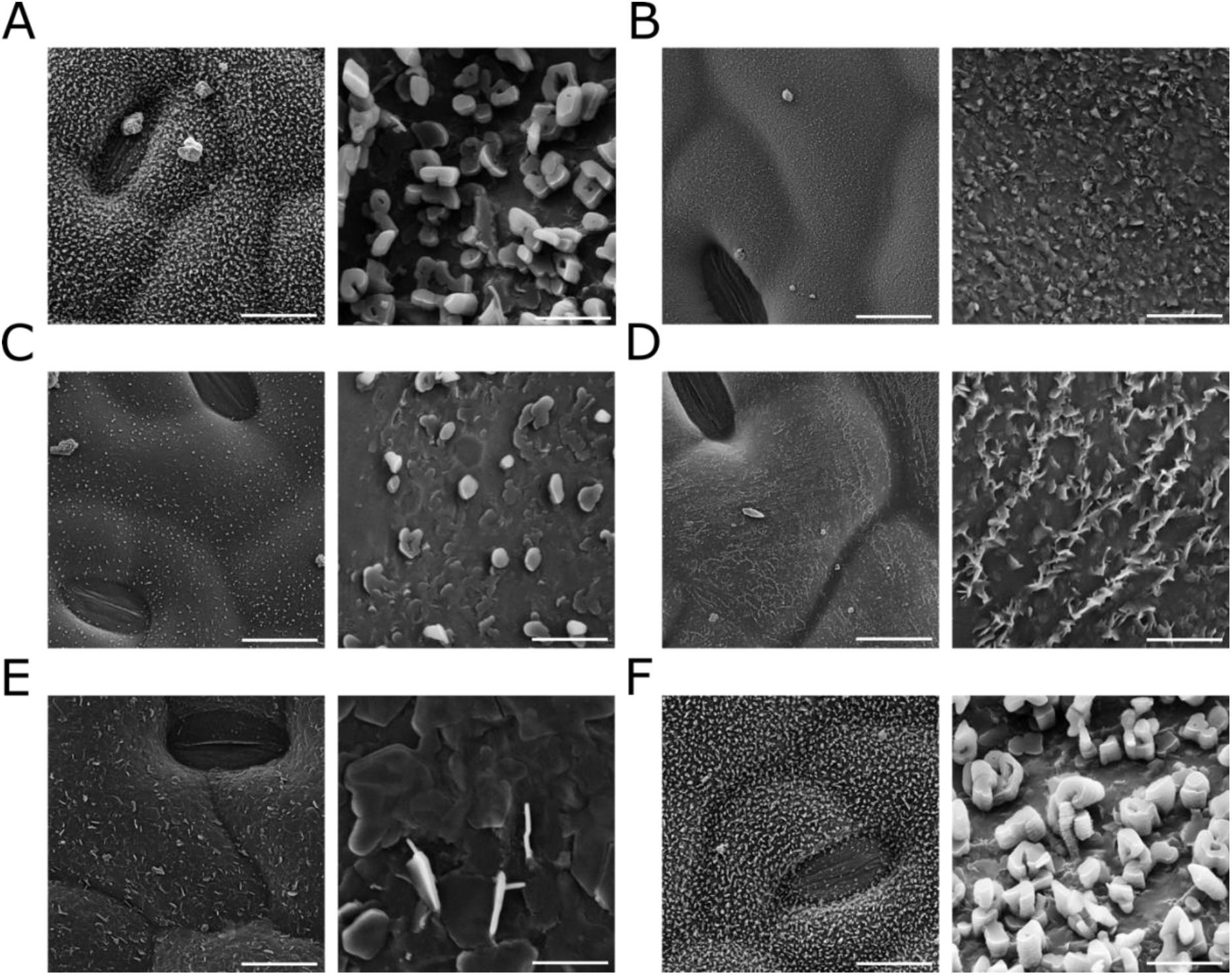
Cryo SEM images of wax crystal dispersion and structure on the adaxial side of mutant plants. **(A)** WT leaf; **(B)** *cer1* leaf; **(C)** *cer3* leaf; **(D)** *kcs6* leaf; **(E)** *far* leaf; **(F)** *abcg32* leaf. For each line, a magnification of x2500 with a 20μM bar is shown on the left and one magnified to x25,000 (2μM bar) on the right.

### Cutin composition of wax metabolism mutants

Wax metabolism and cutin metabolism share some common enzymes and precursors. We therefore examined cutin composition in the same lines analyzed for epicuticular wax (Fig. 6). Here the impact of mutations was far less drastic as compared to their effect on wax composition – not coming close to a reduction percentage of the stronger wax mutants. This was not the case for the *cer1* lines showing significantly altered loads of several cutin monomers, and reaching a reduction of ~64% in methyl caffeate abundance (Fig. 6A-H). Furthermore, when we analyzed the data per chemical class, we found that *cer1* mutants had significantly lower fatty acid load in two out of three lines (Fig. 6B), but a significantly higher levels of ω-hydroxylated fatty acids (Fig. 6C). Mirroring their wax composition, all three *cer1* lines had significantly higher content of very long chain fatty alcohols. Interestingly, *cer1* lines also had reduced total phenol content in their cutin (Fig. 6E).

**Figure 6.**
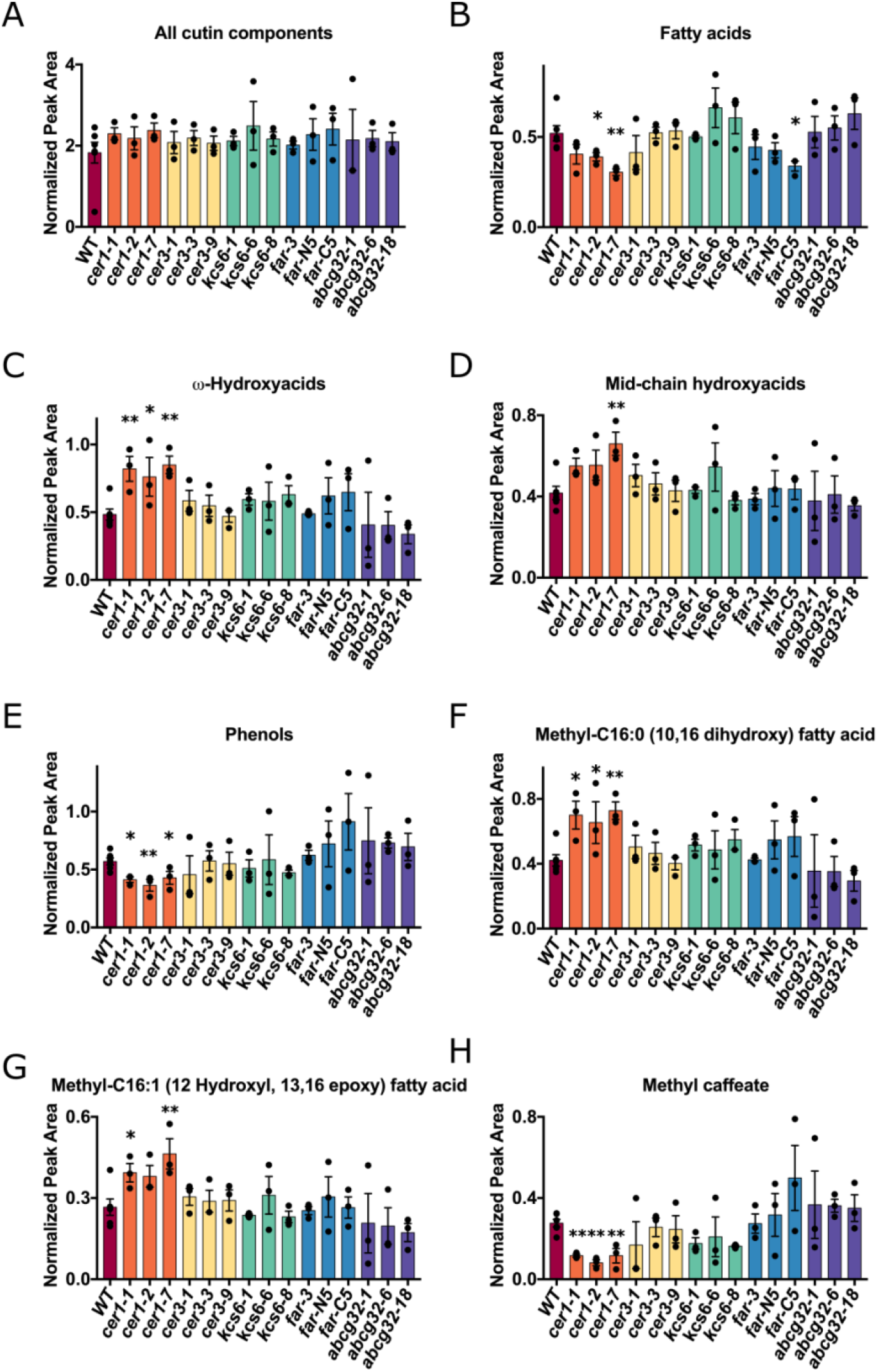
Relative abundance of cutin and its monomers in mutant lines. **(A-E)** Analysis of main cutin monomer classes. **(F-H)** Analysis of most abundant cutin monomers. WT, n=6; all other lines n=3. *= p<0.05 and **= p<0.01 as determined in a student’s t test. Bars represent standard error. All samples were quantified using GS-MS following delipidation, transesterification and derivatization.

### Reduction in alkane abundance increases cuticular water loss drastically

We next examined how changes in chemical composition affected the physiological state of mutant plants by analyzing cuticular water loss. Leaves were detached from plants and weighed every two hours once reaching the linear weight loss stage. Out of the five mutant genotypes, the *cer1* mutants were most strongly affected, showing a three times higher water loss rate (Fig. 7A). The *kcs6* mutant leaves also showed a higher water loss rate whereas *abcg32* leaves displayed a slight and yet significant increase. Surprisingly, of the *cer3* mutant lines only *cer3-1* had a significantly elevated water loss rate. In contrast, two of three *far* lines lost water significantly slower as compared to WT leaves.

**Figure 7.**
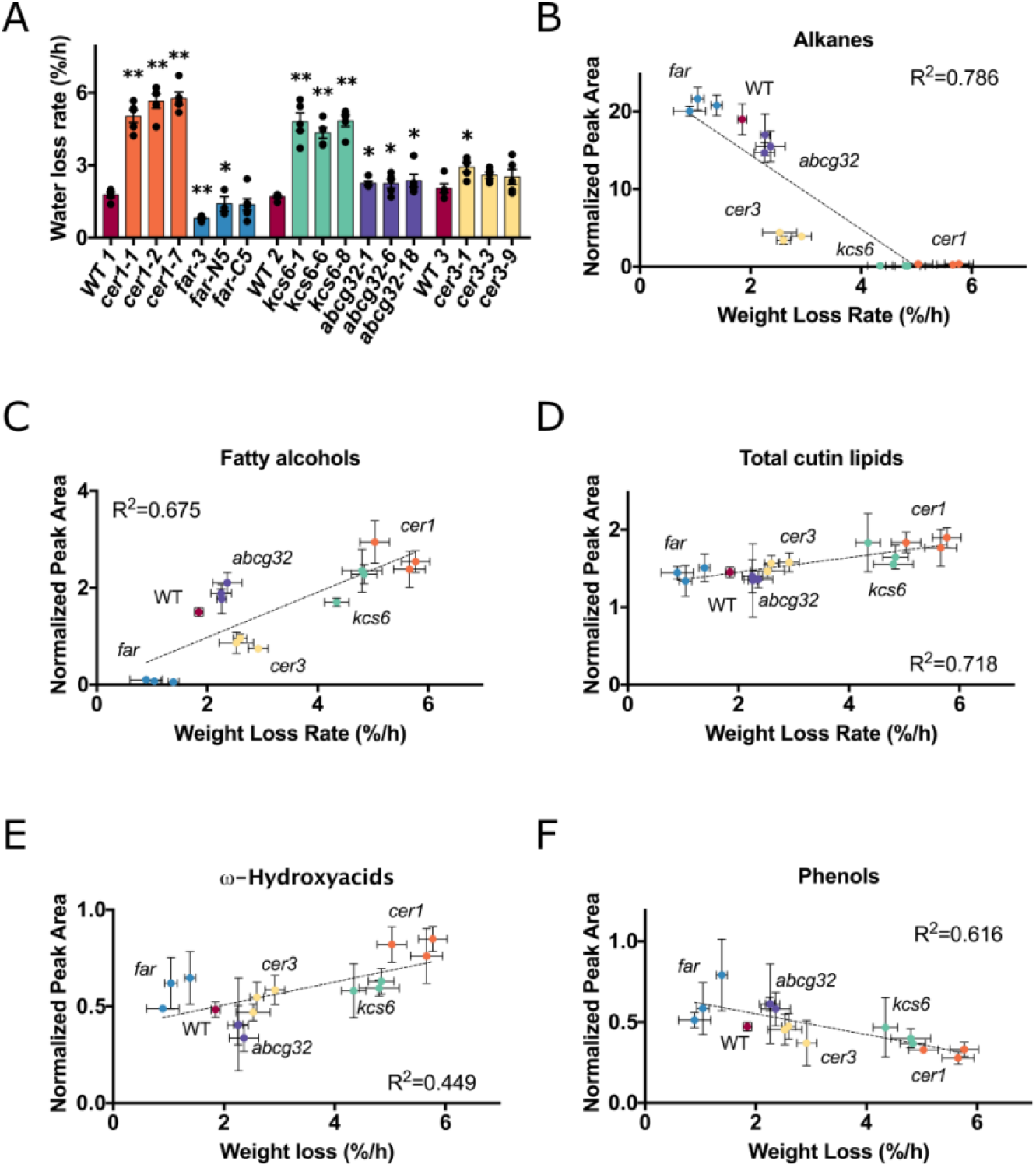
Cuticular water loss rate and its correlation to different wax and cutin components. Cuticular water loss rate. The experiment included three assays in consecutive days each with its own WT samples (WT1 to WT3). Mutant alleles of the same gene are indicated in the same color. * = p<0.05; ** = p<0.01 determined in a student’s t test; n=5. Bars represent standard error. Correlation between water loss rate and **(B)** sum of all alkane wax components, **(C)** sum of very long chain fatty alcohols, **(D)** sum of all lipidic components of the cutin monomers; **(E)** sum of ω-hydroxylated fatty acids of the cutin monomers; **(F)** sum of all phenols of the cutin monomers. R^2^ indicated in each correlation graph is generated from average water loss rate and average abundance of the different wax and cutin components in in the same independent mutant line, obtained in separate experiments.

Next, we plotted water loss rate against the average abundance of wax and cutin components in independent mutant lines (Fig. 7B-F). We found a correlation with an R^2^ of 0.786 between water loss rate and total alkane content. Yet, *cer3* leaves lost water at a slower rate than their alkane content would predict while *cer1* plants lost water at a faster rate than that predicted by the linear regression (Fig. 7B). When cutin components were plotted against water loss, a positive correlation (R^2^ = 0.718) was found with total cutin lipids, although the differences in the cutin component abundance was far smaller between the different lines compared to those in wax components (Fig. 7D). In contrast, phenols found in the cutin fraction (R^2^ = 0.616; Fig. 7F), and especially methyl caffeate (R2 = 0.7) were negatively correlated with water loss. Although the role of the phenolics is not clear to us, these results highlight that alkanes in the epicuticular wax fraction play an important role in preventing cuticular water loss.

### Wax has little effect on water loss pre-drought, but alkanes are essential for recovery from drought

The major effect on cuticular water loss rates due to changes in epicuticular waxes led us to hypothesize that under well-watered conditions plants with greatly reduced alkane abundance would transpire at higher rates and be more susceptible to drought. We tested this hypothesis using a weighing lysimeter system, in which plants are weighed every three minutes and in combination with environmental measurements (e.g. temperature, humidity and light intensity) performed in parallel, transpiration, stomatal conductance (gs), biomass accumulation, water use efficiency (WUE) and drought response can be extrapolated. The first two experiments performed with this system in the summer and winter seasons, involved two and three different mutant alleles of *cer3*, respectively. To our surprise, in the first experiment (during the summer), *cer3* plants had an almost identical daily transpiration compared to WT ones (Fig. S11A). In the ‘winter’ experiment, we observed a trend of *cer3* transpiration starting to be reduced compared to WT as plants grew and their transpiration rate increased, although this trend did not reach significance prior to the onset of drought (Fig. S12A). Furthermore, *cer3* biomass accumulation was not impaired in both experiments (Fig. S11g, S12G), as was their WUE (Fig. S11J, S12I). Plants that respond to drying soil by closing their stomata at a higher soil water content (“theta point”) are more likely to show drought tolerance. Drought response in these two experiments was therefore measured by evaluating this theta point. In the ‘summer’ experiment, *cer3* lines had a theta point similar to WT (Fig. S11K), whereas in the ‘winter’ experiment, their theta point was at a significantly higher soil water content (Fig. S12J). The results indicated that *cer3* plants (displaying approximately 80% reduction in C_31_ alkane) were not negatively affected by their reduced wax load both under well-watered and drought conditions.

The findings described above prompted us to examine whether plants mutated in additional genes that severely affect wax composition would behave in a similar manner. To answer this question, we performed an additional experiment during the spring. Following the previous experiments, we reasoned that during this season we could enjoy the advantages of both the ‘summer’ experiment in which plants grow at a fast rate and the vapor pressure deficit (VPD) and light intensity are relatively stable and the ‘winter’ experiment in which plants are less susceptible to diseases. In this ‘spring’ experiment, we placed three different mutant alleles of either *cer1*, *kcs6* or *far* on the lysimetric system alongside three lines of WT plants. Similar to the *cer3* results under well-watered conditions there was no significant difference between the mutants and WT plants in daily transpiration (Fig. 8A), transpiration rate (Fig. 8D) plant growth rate (Fig. 8G), and water use efficiency (Fig. 8J).

**Figure 8.**
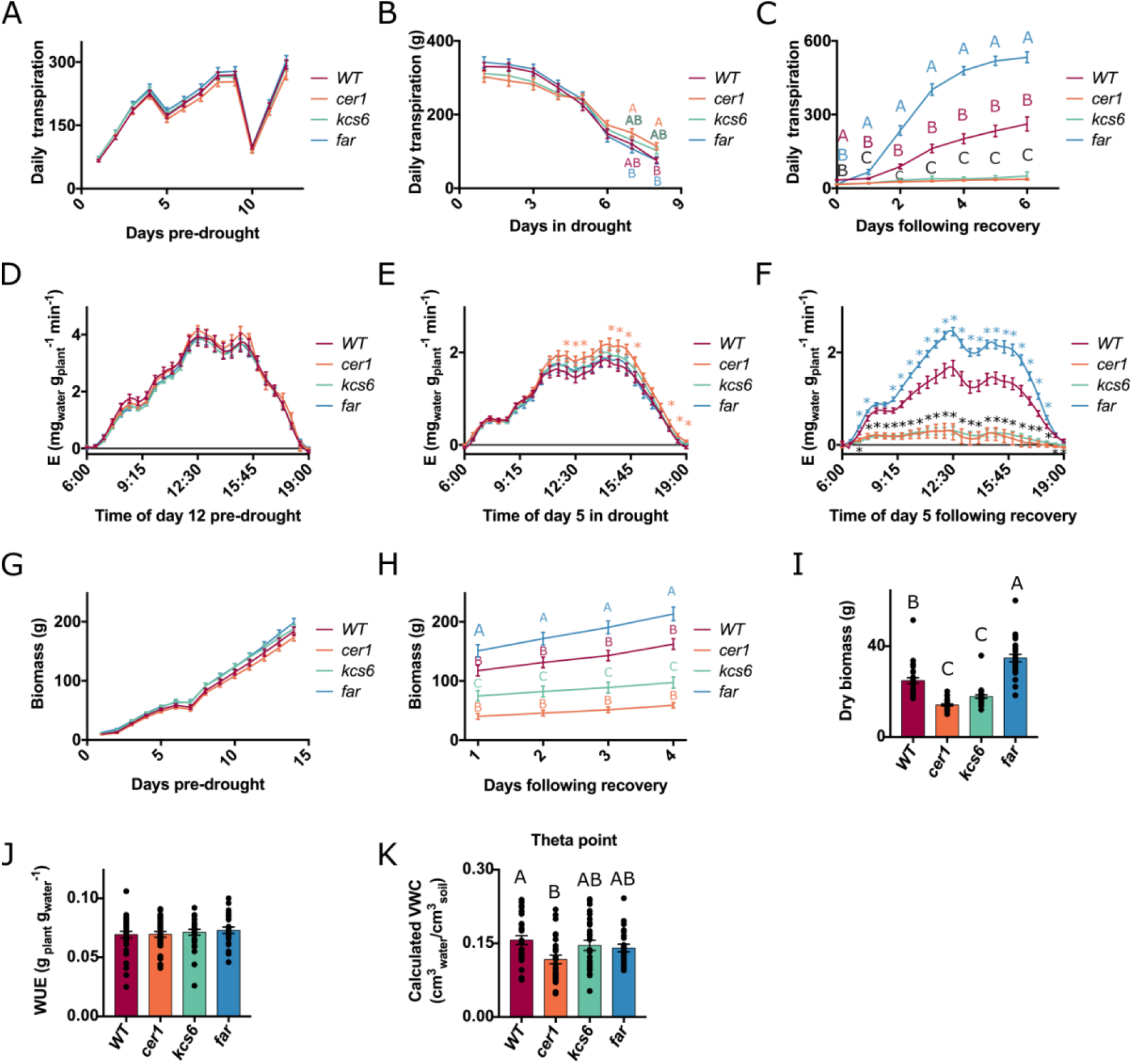
Multi-parameter analysis of mutant plants assayed in a lysimeteric system and assayed following full irrigation, drought and recovery. **(A)** Pre-drought daily transpiration. Daily transpiration following drought initiation and prior to resuscitation. **(C)** Daily transpiration following recovery. Since each individual plant received full irrigation three days after it transpired below the 20% daily transpiration threshold (see materials and methods), the days in recovery unlike those pre-drought and during drought are different for each individual plant, and are averaged according to the number of days past irrigation renewal per plant. **(D)** Transpiration rate during day 12 prior to drought induction. Measurements were taken every 3 min, though error bars are only shown every 30 min. **(E)** Transpiration rate during the fifth day of drought. **(F)** Transpiration rate during the fifth day after plant resuscitation, averaged from different dates for each plant according to its date of recovery. **(G)** Plant biomass during the period prior to drought induction. **(H)** Plant biomass during the four days before the experiment ended, at which time all plants had been resuscitated. **(I)** Dry shoot biomass at the experiments end. **(J)** WUE as calculated pre-drought. **(K)** “Theta point” volumetric water content at which plants began reducing their transpiration rate in response to drying soil. All data is the average of three independent mutant alleles used in this experiment, including WT which had three independent lines grown first in tissue culture prior to the transfer to the greenhouse as performed with all mutant lines. The following amount of lines were analyzed: WT n=28 to 33; *cer1* n=27 to 31; *kcs6* n=25 to 29; *far* n=24 to 26. Different letters indicate significance of p<0.05 as determined in a Tukey HSD test. Asterisks represent significance of p<0.05 as determined in a student’s t test. Asterisks were placed above standard error lines (bars), though points are significant for every point measured between the bars.

Once we exposed well-watered plants to drought and recovery (i.e. re-watering), we detected a markedly different response in the wax mutants. The *cer1* mutant lines’ theta point was significantly lower than WT, while *kcs6* and *far* were similar to WT (Fig. 8K). This led to *cer1* plants having a significantly higher transpiration rate throughout long periods of the day, that was not seen in the other mutant lines (Fig. 8E). Though the increase in *cer1’s* transpiration rate is significant, the effect size is relatively small, and reached a ~20% increase in transpiration rate during the afternoon hours. This was not the case following resuscitation. At this phase, the daily transpiration of *far* plants was significantly higher as compared to WT plants that exhibited significantly greater transpiration than *cer1* and *kcs6* (Fig. 8C, F). These differences rose with time up to the sixth day following resuming irrigation; *far* plants transpired 533ml a day on average, WT 262 ml, *kcs6* 50 ml and *cer1* transpired only 36 ml on average during that day (Fig. 8C). These results were mirrored by the transpiration rate (Fig. 8F), plant growth rate (Fig. 8H) and shoot dry biomass (when the experiment ended; Fig. 8I). The underlying reason for these drastic differences was evident when we examined the plants once completing the experiments (Fig. S13). While all *far* plants retained their original leaves, some WT plants were left with leaves and others not. In a vast majority of the *cer1* and *kcs6* plants leaves present at the time of drought initiation dried and the transpiration was from new leaves, which started growing from axillary buds upon resuscitation (Fig. S13).

### Drought followed by resuscitation leads to stem cracking

Towards the completion of the weighing lysimeters experiment, we detected an intriguing phenotype, which to the best of our knowledge has not been reported in the context of wax deficiency. Sixteen-out of 27 *kcs6* lines derived from all three independent mutant alleles displayed severe cracks along their stems penetrating the xylem and deep into the stem pith (Fig. 9A-B). Intriguingly, these cracks even led to water dripping freely out of the xylem and out of the cracks (Fig. 9A). Besides the *kcs6* plants, there was one occurrence of stem cracking in a *cer1-1* plant indicating that the occurrence in *kcs6* plants was not an isolated event. When investigating the reasons for the *kcs6* stem phenotype we found that the 11 *kcs6* plants with intact stems whose non-cracking phenotype was not due to technical reasons had a similar biomass and theta point compared to the plants with cracked stems at the onset of drought (Fig. 9E; Fig. 9F). While all 16 cracked plants resuscitated during three days, from the 13^th^ day of drought to the 15^th^ day, the uncracked plants were resuscitated between the 10^th^ to the 18^th^ day in drought. When examining the 11 *kcs6* plants with undamaged stems we found that they reduced their volumetric water content (VWC) at a slower rate than those with cracked stems, though this difference was not significant on a point measurement basis (Fig. 9G). Thus, as is seen from the wider range of drought duration prior to recovery, plants that depleted their water and resuscitated rapidly as well as those that transpired less and were exposed to drought more gradually did not experience stem cracking.

**Figure 9.**
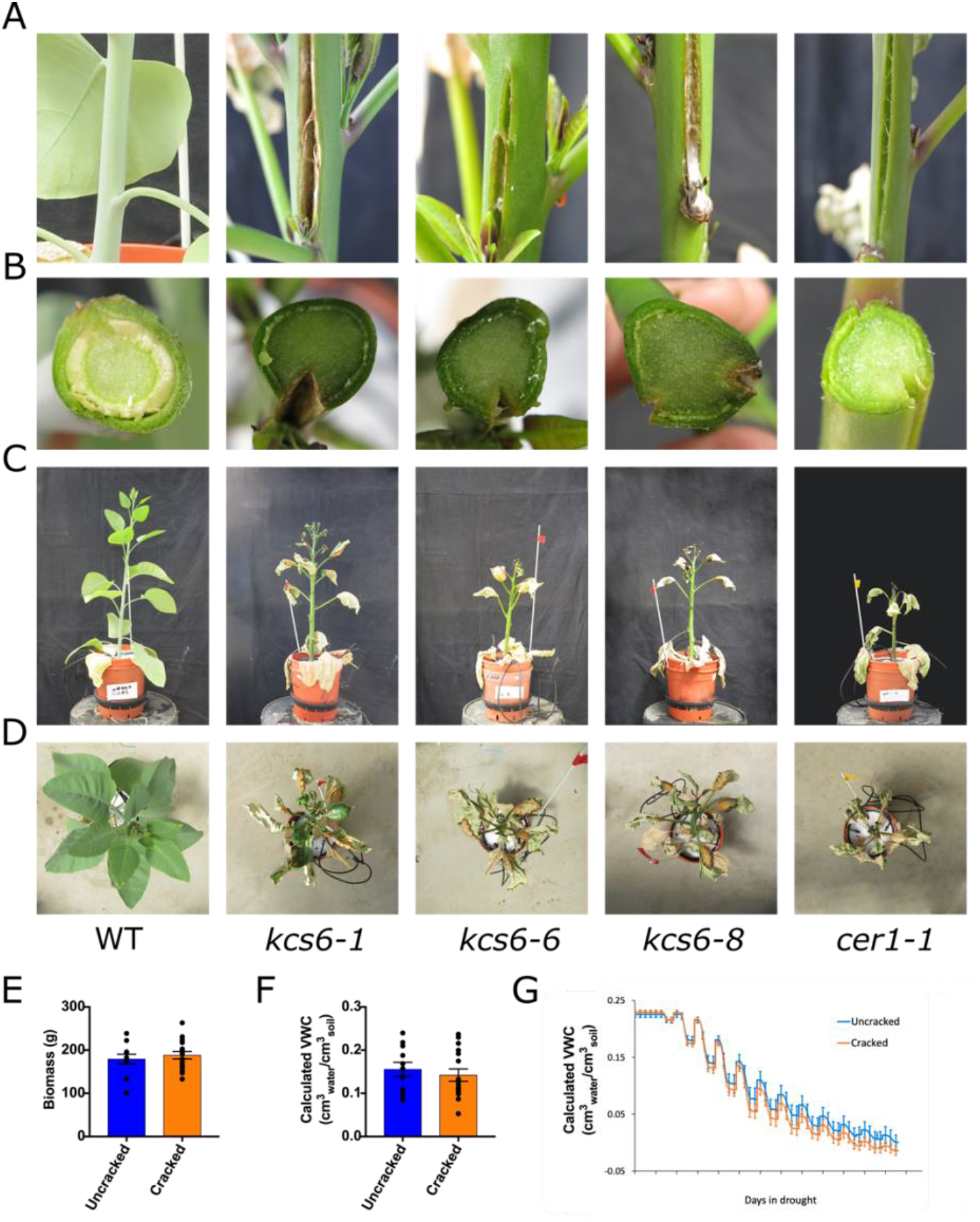
The stem cracking phenomenon. **(A)** Images of cracked stems phenotypes in representative plants from the three *kcs6* independent mutant alleles, a *cer1* plant displaying a similar phenotype and a representative WT stem. **(B)** Images of cut stems showing the radial formation of the crack through the xylem and into the stem pith cells. **(C)** Images of whole plants at the experiments end. **(D)** Whole plants photographed from above and displaying differential drought response. **(E)** Biomass of plants with or without cracked stems (*kcs6*) measured during the last irrigated day prior to drought. **(F)** Volumetric water content of *kcs6* plants in which no stem cracking was observed compared to those in which stems cracked. Plants included in this analysis were of similar size at drought onset. **(G)** VWC in pots of cracked and uncracked *kcs6* plants as drought commenced. Cracked n=16; uncracked n=11.

### Stomatal aperture and development do not compensate for elevated cuticular water loss

We examined whether the discrepancy between high leaves cuticular water loss and normal transpiration rate under well-watered conditions was related to a reduced stomatal density, size or aperture, leading to the elevated cuticular water loss being compensated for by reduced stomatal water loss. However, all mutants had similar stomatal density (Fig. S14A, D) and sizes (Fig. S14B,E) compared to WT, and even those insignificant differences in density were compensated for by larger stomata; i.e. plants with a trend of less dense stomata had a trend of larger stomata as well. Abaxial stomatal aperture (Fig. S14C) was the only parameter where we found a significant difference, with *cer1* having a larger stomatal aperture. Thus, unlike our original hypothesis, cuticular water loss was not compensated for by stomatal aperture or morphology.

### The glossy phenotype of wax mutants does not significantly affect light response and plant fitness

Despite extreme differences in their wax composition, *cer1*, *cer3*, *kcs6* and *far* mutants, all displayed a glossy leaf appearance. To examine if this phenotype affects photosynthetic efficiency, we examined RUBISCO efficiency (Vc_max_) and maximal electron transport rate (J_max_) by monitoring carbon assimilation as sub-stomatal CO_2_ concentration was raised (A/Ci curves). The *cer1* mutant lines appeared different from all other mutants and the WT as their A/Ci curve plateaued much earlier than all other genotypes (Fig. 10A). Although the J_max_ of *cer1* plants was not significantly lower (Fig. 10C), their Vc_max_ was significantly reduced (Fig. 10B). *cer1’s* wax composition is similar to *kcs6*, a fact that leads to very similar responses in assays where wax composition is the underlying factor. However, *cer1* is the only mutant with altered cutin, leading us to suggest that the differences seen in the A/Ci curves were the result of altered cutin and not wax load or composition.

**Figure 10.**
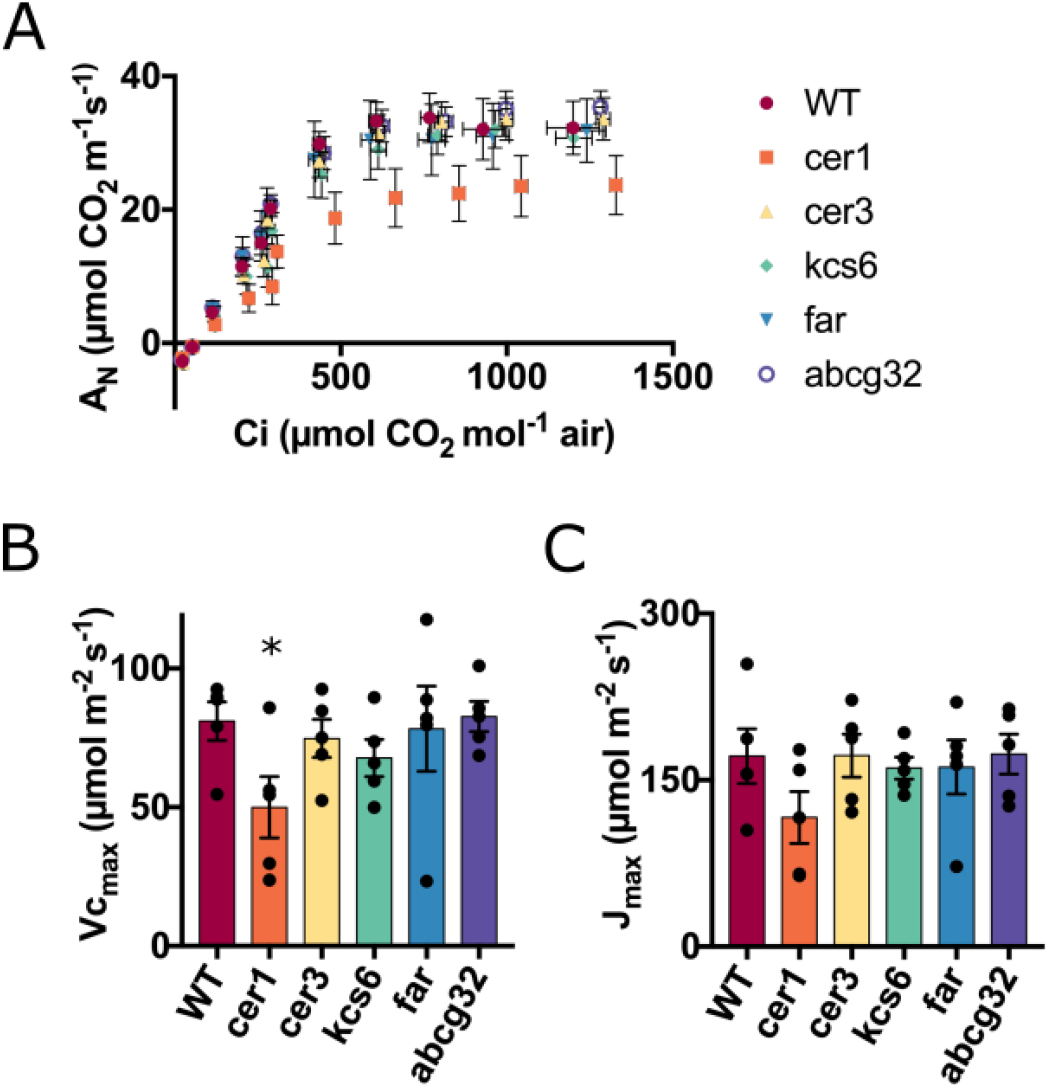
Photosynthetic efficiency of WT and four wax biosynthesis mutant lines as analyzed by A/Ci curves **(A)** Average carbon assimilation with rising Ci concentrations. **(B)** Vc_max_ indicating RUBISCO efficiency. **(C)** J_max_ indicating maximal electron transport rate. Asterisks indicate a significance of p<0.05 as determined in a student’s t test. N=5.

We next asked if the glossy phenotypes of wax mutants had a significant impact on light response and consequently plant fitness. Thus, we exposed plants of each genotype to 12 rising light intensities and recorded carbon assimilation and stomatal conductance (Fig. 11A and Fig. 11B). No plants reached photoinhibition along these points despite reaching 2200 μE (while plants were grown at a light intensity of 100μE – 200 μE). Similar to the A/Ci curves results, *cer1* mutants were the only plants displaying an altered phenotype exhibiting reduced carbon assimilation at five points of the dozen light intensities measured (Fig. 11A). These results demonstrate that light reflectance by wax crystals does not alter photosynthetic capacity under non-stressed conditions.

**Figure 11.**
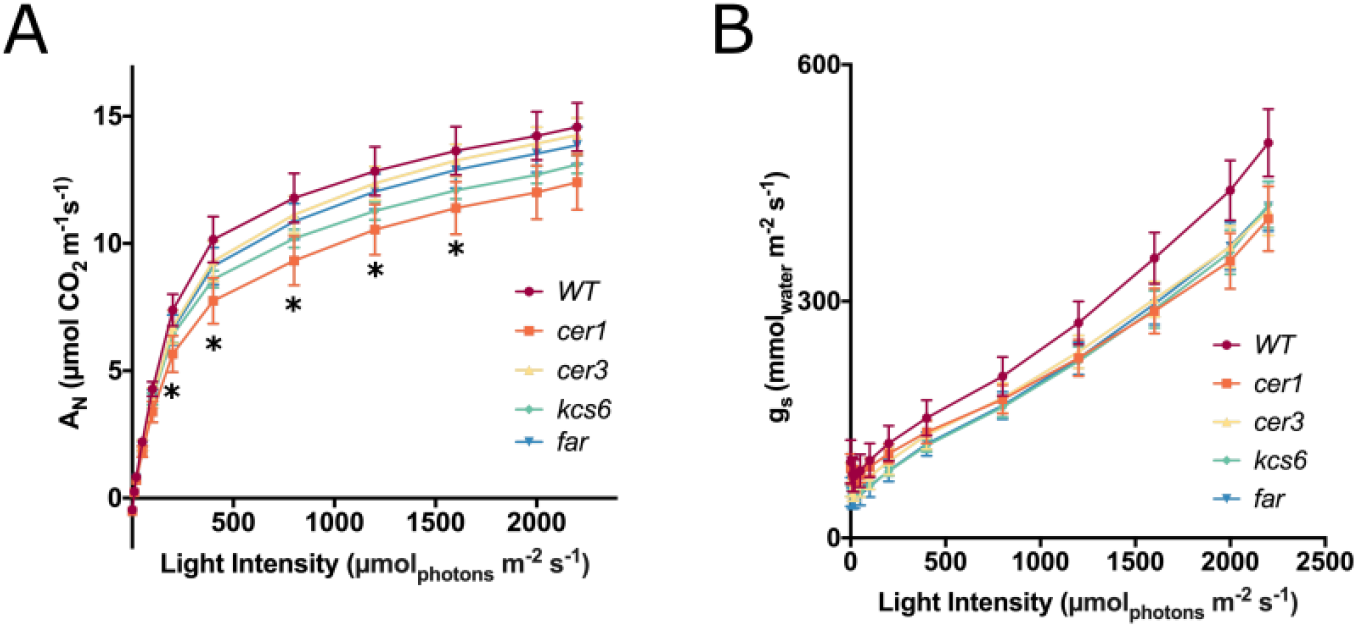
Effects of wax deficiency on gas exchange in response to rising light intensity. **(A)** Carbon assimilation. **)B(** Stomatal conductance. Asterisks indicate a significance of p<0.05 as determined in a student’s t test. N=9.

## Discussion

### Tree tobacco in cuticular lipids research

For many years, cuticular lipids research has been largely performed in *Arabidopsis thaliana* (Lee and Suh, 2015). The ease of screening large mutant collections enabled identification of the majority of *eceriferum* mutants and their characterization. Advancements in targeted mutagenesis, predominantly the CRISPR-Cas technology, opened the way for exploiting other species such as tree tobacco in cuticular lipids research. Apart from outstanding advantages including an extremely high wax load, simplicity of wax composition and induced wax accumulation following drought, tree tobacco is readily transformed, can grow in a variety of conditions including natural settings, and recently had its genome sequenced (Usadel et al., 2018). Furthermore, being a perennial plant, tree tobacco can be explored at both the juvenility stage as well as upon maturation and bark formation. Hence, several key results reported here could not have been discovered in Arabidopsis, such as stem cracking and drought experiments in which plants accumulated tens of grams of biomass and transpired hundreds of milliliters each day.

### Alkanes reduce cuticular water loss, but cutin affects photosynthetic sufficiency and light response

In this study, we simultaneously characterized a large number of mutants corresponding to different cuticular lipids genes. By doing so, we filtered out indirect effects not related to epicuticular wax composition such as changes in cutin composition. Both *cer1* and the *kcs6* mutants possess an extremely reduced alkane load, yet, *cer1* also retains an altered cutin composition, with significantly less phenols and more ω-hydroxylated fatty acids. Comparing the phenotype of *cer1* to that of *kcs6* enabled us to make the distinction. Cuticular water loss is affected when both genes are mutated, although the rate in *cer1* is higher. Here we detected an additive effect of altered cutin despite the major effect being that of alkane abundance. Effects that are only present in *cer1* and not *kcs6* and which we attribute to cutin rather than epicuticular wax are the reduced Vc_max_ and lower carbon assimilation with rising light intensity. In contrast to these, drought response in *cer1* and *kcs6* plants was nearly identical, suggesting that alkane abundance is a major factor essential for recovery following drought.

It was previously shown that overall accumulation of epicuticular waxes following exposure to drought in tree tobacco reduced the rate of cuticular water loss (Cameron et al., 2006). However, Cameron et al. could not determine which of the many wax constituents underlays this reduction in water loss rate. Here we point to alkanes being the component preventing cuticular water loss. While the correlation of alkanes with water loss rate is high, it does not seem to be linear. The *cer3* mutants possess ~20% of the WT C_31_ alkane and had only a slight increase in water loss rate. This discrepancy could be observed when removing the *cer3* lines from the regression, which raises the R^2^ from 0.79 to an 0.95 value. It therefore seems that although alkanes reduce cuticular water loss and are essential for recovery following drought, the threshold amount of these components required for having an effect may be well under the one present in WT plants.

### Alkane abundance is essential for recovery following drought but does not contribute to fitness under well-watered conditions

Following the results of the cuticular water loss assay performed on detached leaves, we anticipated that assaying whole plants would mirror this at least partially, and that the mutant plants would transpire at a higher rate under well-watered conditions. This is in line with many studies reporting epicuticular wax conferring drought tolerance, achieved in many cases due to reduced transpiration (reviewed in Xue et al., 2017). Our conclusions from three independent whole plant growth and drought experiments do not match the common view point that wax simply confers drought tolerance by reducing transpiration rates. Plants from our mutant collection, regardless of the metabolic impact of the mutation (whether having no fatty alcohols, no alkanes or a reduced alkane load) all transpire and accumulate biomass at a rate similar to WT under well-watered conditions. Furthermore, although *cer1* responded to drought by reducing stomatal conductance at a lower soil water content (a phenotype associated with reduced drought tolerance; Negin and Moshelion, 2017), such a response was not detected in *kcs6* plants. However, this was not the case with the mutants’ recovery dynamics, in which extreme differences could be seen between plants, clustering according to their wax composition. We found that the underlying phenotype that determined whether plants would recover transpiration and growth rapidly following irrigation resumption was leaf survival. We observed clearly that plants that did not recover, were lines in which the drought caused leaves to dry out and die (Fig. S13). *Far* mutant plants in contrast to alkane reduced ones, recovered at the fastest rate, and lost almost no leaves during the drought treatment.

In our drought assay, soil water content and transpiration were continually monitored and plants were resuscitated once they had reached the same transpirational threshold. In contrast, many studies assay drought by stopping irrigation for a similar duration of time (see for example Bourdenx et al., 2011). This experimental design causes plants that have higher transpiration under well-watered conditions to reach drought conditions at an earlier time, and be exposed to drought conditions for a longer period than their counterparts. Furthermore, “drought tolerance” attributed to altered drought response may in fact be secondary to attenuation of other stress conditions. These stress conditions may include high light intensity, temperature and VPD. For example, plants that are able to disperse light rather than absorb it may have a lower leaf temperature, which may be compensated for by increasing transpiration (Richards et al., 1986), making these plants less tolerant to drought induced by a similar duration of irrigation stoppage. Our experimental setup uncoupled well-watered transpiration and growth from drought response and recovery. This enabled us to point to the ability to strongly reduce transpiration under drought and prevent leaf death as alkane dependent and essential for recovery.

### Changes in wax composition and the stem cracking phenomena

The phenomena of stem cracking is known in conifers at the end of growing seasons of drought years, and increases with drought severity (Zeltińš et al., 2018; Cameron, 2019). However, to the best of our knowledge it has never been linked to alterations in epicuticular wax abundance or composition. Its appearance in *kcs6* and *cer1* mutants following recovery indicates that stem cracking is indeed alkane related. The extensive use of annual plants, mainly Arabidopsis for molecular research of epicuticular waxes, has left its effect on perennial and woody plants relatively unexplored. Apart from presenting an intriguing phenotype linked to deficiencies in epicuticular wax biosynthesis, our study here underscores the use of *N. Glauca* as a model system enabling research of wax function in woody species. The exact stage at which stem cracking took place is not clear to us. Even at the end of the drought treatment, we could not detect cracked stems in the *kcs6* plants. However, whether small initial cracking already began during drought, if a more gradual recovery would have prevented the cracking, and what is the threshold of drought conditions resulting in cracking, remains to be determined.

### Glossy mutants did not share a common phenotype under the examined conditions

Out of the five mutants examined in this study, four display a glossy phenotype. This striking phenotype appears in plants that lost alkane biosynthesis nearly entirely but also when the C_31_ alkane is greatly reduced in *cer3* mutants, and even in the *far* mutants that are deficient merely in primary alcohols. The *far* mutants do not exhibit an elevated rate of cuticular water loss. More strikingly, their recovery following drought is greatly improved compared to WT indicating that fatty alcohols are completely unnecessary for drought response. This leads to the question of what is the disadvantage in losing primary alcohols specifically and exhibiting a glossy appearance? Since the glossy phenotype is dependent on optical parameters, we examined photosynthetic efficiency as well as light response. In both cases, there was no correlation between leaf glossiness and these parameters. In eucalyptus species, it was shown that glaucous leaves had the effect of reducing photosynthesis prior to saturating conditions. In addition, in these plants, once wax was removed, photoinhibition occurred at high light intensity while this did not occur in normal plants (Cameron, 1970). Although our initial hypothesis was similar to the findings of Cameron both in terms of reaching photoinhibition at a lower light intensity as well as having higher carbon assimilation at a non-saturating light intensity, this was not the case in the four mutants examined in this study. The *cer1* mutant line even assimilated carbon at lower rates with rising light intensity.

Epicuticular wax has been suggested to play a protective role against a range of stresses. Drought (Aharoni et al., 2004; Seo et al., 2011; Lee et al., 2014; Lee and Suh, 2015; Xue et al., 2017), UV radiation (Long et al., 2003), osmotic stress (Liu et al., 2019) and insect herbivory (reviewed in Eigenbrode and Espelie, 1995) are a few such conditions. In addition, since epicuticular wax repels water it has been suggested to have an anti-adhesive self-cleaning affect (also known as the “lotus effect”; Neinhuis and Barthlott, 1997), which causes different particles including pathogens to be washed away upon leaf wetting. However, under our experimental conditions, we did not find a common disadvantage in coping with abiotic stress conditions which is linked to a glossy appearance. Hence, we suggest that a glaucous appearance is likely to bear positive effect in combating biotic stress conditions rather than under abiotic ones.

### Conclusions

The examination of epicuticular wax mutants in *N. glauca* revealed that while alkane accumulation is strongly correlated with cuticular water loss, under well-watered conditions epicuticular wax composition does not affect whole plant water loss or plant growth rate. Despite this, the presence of alkanes greatly affected the ability of plants to recover following drought conditions, and its deficiency led to a previously unobserved phenotype of stem cracking. Furthermore, under our experimental conditions, we could not find a common denominator between plants possessing a glossy phenotype suggesting that a glaucous appearance is associated with combating biotic stress conditions. In contrast to this, plants in which cutin was strongly affected were the only ones whose photosynthetic efficiency and light response were altered, emphasizing the importance of uncoupling cutin related affects from those which are epicuticular wax derived.

## Materials and methods

### Plant material and growth conditions

Seeds of *N. glauca* were initially collected from plants growing nearby the Weizmann institute campus in Rehovot, Israel (31.912408, 34.820184). Plants used for transcriptome analysis were grown in the greenhouse (winter 2013). Plants used in cuticular wax and cutin extraction, SEM, stomatal analysis, cuticular water loss and photosynthetic efficiency assays were grown in a greenhouse in two batches during the winter-spring of 2019 and of 2020. Plants used for light curves were grown in a growth room under a flux of ~150μE, with a temp. of 22°C and light/ dark period of 16/8 h. Drought experiments were performed in the lysimetric facility in the Faculty of Agriculture in Rehovot, Israel (31.904134, 34.801060), during 7-9.2018, 1-3.2019 and 3-4.2020.

### CRISPR vectors and mutation analysis

Construct assembly was performed using the ‘golden braid’ cloning system (Sarrion-Perdigones et al., 2013). A codon optimized *Streptococcus pyogenes* Cas9 (Fauser et al., 2014) was used, driven by the *Solanum lycopersicum* UBIQUITIN10 promoter (Dahan-Meir et al., 2018). All gRNA transcription was driven by the Arabidopsis U6-26 promoter. crRNAs were planned based on the genome assembly of Usadel et al. (2018) using the Crispr-p software (Lei et al., 2014). crRNA sequences, gene maps and transformation protocol appear in supplementary methods and Fig. S2-S10. Mutations were analyzed by DNA extraction and PCR amplification using oligonucleotides flanking the expected edited region. In plants where the crRNAs were positioned far from each other on the genome, two PCRs with different primer sets were performed and PCR products sequenced. For homozygous mutations, NCBI blast (https://blast.ncbi.nlm.nih.gov/Blast.cgi) was used to compare the mutant’s sequence to WT sequence and in heterozygous cases, they were characterized using one of two softwares: DSDecode (Liu et al., 2015) or CRISP-ID (Dehairs et al., 2016).

### Transcriptomics and differential gene analysis

Transcriptome analysis was performed on plants from several tissues under well-watered and drought conditions [induced by three events of drying and recovery as described in Cameron et al. (2006)]. RNA extraction, library preparation and RNA-seq were performed as described in Hen-Avivi et al. (2016). Since at the time of analysis of the raw reads, the *N. glauca* genome was unpublished, we performed de-novo transcriptome assembly based on Haas et al. (2013) of *N. glauca* gene expression data published by Long et al. (2016). Based on this assembly, genes were aligned and their differential expression analyzed. This data was then used to define conditions of epidermis enrichment and drought induction, and contigs in which at least three conditions (adaxial epidermis enrichment, abaxial epidermis enrichment and adaxial epidermis drought induction for example) were searched using BlastX against the NCBI protein sequence database (NR) to find their closest homologs.

### Wax and cutin monomer profiling

For wax extraction, three leaf discs with a diameter of 12mm each were dipped in 4ml chloroform with an internal standard of 10μg C_36_ alkane and shaken gently for 15sec. The discs were then removed and chloroform was evaporated under a nitrogen flow. Samples were then resuspended in 100μl chloroform to which 20μL pyridine and 20μL BSTFA were added. Samples were derivatized at 70°C and injected in a splitless mode to a GC-MS system (Agilent 7890A chromatograph, 5975C mass spectrometer and a 7683 auto sampler) as described in Cohen et al. (2019). Chromatograms and mass spectra were analyzed using MSD Chemstation software (Agilent). Identification was based both on fragmentation alignment to the NIST Mass Spectral Library and in-house retention Indexes. Quantification was performed using the Chemstation software and was normalized to the C_36_ alkane internal standard. Cutin extraction was performed as described in Cohen et al. (2019). Monomer identification and quantification was performed as in wax components analysis, with normalization performed against a C_32_ alkane internal standard.

### Leaf desiccation assays and whole plant drought trials

Five leaves from different plants from each independent line were cut in the greenhouse and immediately inserted to zip-locked bags, brought to the lab and let dry at 22°C. Leaves were weighed every two hours for 12 hours. This was done following an earlier calibration in which leaves were weighed every half hour and which showed that following two hours of desiccation water loss was linear up to 24h. Due to the large number of leaves and need to repeatedly weigh them at given intervals, the experiment was divided to three different days, each having its own WT control. Drought trials performed using a lysimetric system (during the summer of 2018, winter of 2019 and spring of 2020) were performed in either soil (2018, 2019), or sand (2020) as described in Halperin et al. (2016) and Dalal et al. (2020) (see supplementary methods for details).

### Stomatal analysis and scanning electron microscopy

Stomatal imprints were taken from adaxial and abaxial epidermises of greenhouse grown plants, as described in Yaaran et al. (2019). Nail polish imprints of dental resin were then attached to microscope slides and photographed in a Nikon eclipse E800 microscope with a Nikon Digital sight DS-5Mc camera. Each imprint was photographed at two different locations at a magnification of x100 for stomatal density and at another two at a magnification of x400 for stomatal aperture and size. Stomata were then quantified and aperture and size were measured using ImageJ software (https://imagej.nih.gov/ij/). For SEM, small sections were cut from fresh leaves of WT and representative lines from the five mutant genes and inserted to a cryo-holder, frozen in liquid nitrogen, coated and photographed using a Zeiss Ultra 55 SEM as described in Hen-Avivi et al. (2016).

### Photosynthetic efficiency and light response

Photosynthetic efficiency was assessed by measuring gas exchange at rising CO_2_ concentrations (“A/Ci curves”), using a Li-6800 portable photosynthesis system (LI-COR, Inc.; Lincoln, NE, USA). Parameters were set to: PAR 1,600, relative humidity 60%, CO_2_ 400PPM and leaves were inserted to the infrared gas analyzer (IRGA) chamber for acclimation. This stage was performed for at least 15min and was stopped when carbon assimilation ceased rising exponentially (but no longer than 25min). A/Ci curves were then performed with the points: 400PPM, 300PPM, 150PPM, 50PPM, 0, 400PPM, 600PPM, 800PPM, 1,000PPM, 1,200PPM and 1,500PPM CO_2_. Data was analyzed using R (www.r-project.org) and the ‘plantecophys’ package (Duursma, 2015). Light response was measured using the Li-6800 portable photosynthesis system. Parameters were similar to those used for A/Ci curves, excluding light intensity which was raised every 6 min. Plants were first dark acclimated by covering the leaf on which the measurement would be performed in aluminum foil for 10min, after which the leaf was inserted to the IRGA chamber. Each light intensity was kept for at least 6min, and once A and gs were stable a measurement was taken. The light increment program was as follows: 0, 10 μE, 20 μE, 50 μE, 100 μE, 200 μE, 400 μE, 800 μE, 1,200 μE, 1,600 μE, 2,000 μE and 2,200 μE.

### Statistical analysis

The JMP 14 software (SAS Institute; http://www.jmp.com/en_us/home.html) was used for all statistical analyses, except for two-piece linear curves in the drought experiments in which in-house statistical tools of the lysimeter system (https://www.plant-ditech.com/) were used to find the best fitting regression lines. Student’s t test was used when comparing two groups. Mutants were always compared to WT, except in the spring drought trial where Tukey HSD test was used and all groups were compared.

## Acknowledgments

We thank Prof. Björn Usadel for providing us with access to unpublished *N. glauca* genomic and transcriptomic data. These high quality assemblies aided us to a great extent throughout this study.

