## Supplemental table 1 for "Cuticular Wax Composition is Essential for Plant Recovery Following Drought with Little Effect under Optimal Conditions"

Table S1: the 16 genes targeted for knockout mutations in this study, their function and Arabidopsis homologs.

| Gene | Function | Role | Arabidopsis homolog |
|---|---|---|---|
| *NgSHN1-like* | Transcription factor | Regulates wax and cutin synthesis | *AtSHN1* - AT1G15360 |
| *NgSHN3-like* | Transcription factor | Regulates wax and cutin synthesis | *AtSHN3* - AT5G25390 |
| *NgMYB96-like* | Transcription factor | Regulates wax synthesis in response to drought | *AtMYB96* - AT5G62470 |
| *NgGPAT4-like* | Cutin synthesis | Attaches cutin monomers to glycerol | *AtGPAT4* - AT1G01610 |
| *NgCYP86A22-like* | Cutin synthesis | ω-hydroxilates cutin monomers | *AtCYP86A7*- AT1G63710 |
| *NgGDSL-like* | Cutin synthesis | Polymerizes cutin monomers | *AtGDSL* - AT1G75900 |
| *NgLACS-like* | Wax and cutin synthesis | Converts fatty acids to acyl-CoA and vis versa | *AtLACS1* - AT2G47240 |
| *NgKCS6-like* | Wax synthesis | Part of the fatty acid elongase complex | *AtCER6* - AT1G68530 |
| *NgCER1-like* | Wax synthesis | Together with *CER3,* converts acyl-CoA to alkanes | *AtCER1* - AT1G02205 |
| *NgCER3-like* | Wax synthesis | Together with *CER1* converts acyl-CoA to alkanes | *AtCER3* - AT5G57800 |
| *NgMAH1-like* | Wax synthesis | Hydroxylates alkanes to form secondary alcohols and ketones | *AtMAH1* - AT1G57750 |
| *NgFAR-like* | Wax synthesis | Reduces acyl CoA to form primary alcohols | *AtCER4* - AT4G33790 |
| *NgWSD1-like* | Wax synthesis | Combines fatty alcohols and fatty acids to form wax esters | *AtWSD1* - AT4G33790 |
| *NgABCG11-like* | Wax and cutin monomer transport | Transports cutin monomers to the extracellular matrix | *AtABCG11* - AT1G17840 |

| | | | |
|---|---|---|---|
| *NgABCG32-like* | Wax and cutin monomer transport | Involved in cutin transport to the extracellular matrix | *AtABCG32* - AT2G26910 |
| *NgACBP-like* | Wax and cutin monomer transport | Involved in lipid transport | *AtACBP1* - AT5G53470 |
