## Supplemental methods for "Cuticular Wax Composition is Essential for Plant Recovery Following Drought with Little Effect under Optimal Conditions"

### Supplementary methods

*CRISPR constructs assembly and transformation*

The crRNAs used for mutation induction were as follows: *ABCG11*- 5'-CTCAACTACGGCTACCTGAT and 5'-CACTTGCGTGGTATAAGTGG. ABCG32:5'-AATACAGTTCCCCAAGATAT and 5'-GGAAGCGTATAGACCCAGCA. ACBP1: 5'-GTAGTTGATCAGTCTGAAAA and 5'-TACTGCTCCATAGCCACCTC. CER1: 5'-TTCTCACAACTGTGCTAACT and 5'- TGTAAGCTGTGAGTCACATA. *CER3*: 5'-GATGCGGCATGCATCATACC and 5'- TATTATATTAGCTGCAGTGA. *CYP86A22*: 5'-TCAATTCAGATGGTGACACG and 5' – CCGCGATCCATTCATGCATT. *FAR*: 5'-GAAAAGGCCACGATTATAGC 5'- ACTTCAGCCATGAAGGAGCT and 5'-TAGTCCTGATGTACCGTTCA. *GDSL*: 5'GGAGATTCCATTGTTGATCA. *GPAT4*: 5'-TCCGTCGCGGCGGATCTCGA and 5'- GTGAAGCGAGCTCGATATCT. *KCS6*: 5'-AAGTTACAATCTTTCTGGTA and 5'- GGAGGGATGTAATGACAAGC. *LACS*: 5'-CGGAGTGCACACATGCTAAA and 5'- GTGTTTCACTTCGCTGCAGC. *MAH1*: 5' –TTACGAAATTCGGGGCCCTT and 5'- TCGTTGAAGCCCTTTTCACA. *MYB96*: 5'-GCAAAAGCTGTAGACTGCGA and 5'- CTCGAAACCGCACCGATTGA. *SHN1*: 5'-AGGTGTCAGGCAACGCCATT and 5'- CCCTTCTTCTGATTCGAAAT *SHN3*: 5'-AGGTGTCAGACAACGCCAGT and 5'- GATAACGATAATTGTCACAT *WSD1*: 5'-GAAACCAATCGCCGATAGTA and AGAAGTTGCCTCGGCATGAG. For maps of gene structure and target location of the nine genes that displayed phenotypes in T0, see figures S2-S10. Constructs were transferred to the GV3101 *Agrobacterium tumefaciens* strain for plant transformation. Stable transformation was based on the method presented in Mozo et al. (1992). *N. glauca* seeds were surface sterilized in NaClO (3%) for ten minutes, after which seeds were rinsed twice in autoclaved DDW. Seeds were then sown magentas containing Nitsch medium (Duchefa Biochemie). *A. tumefaciens* GV3101 containing the construct of interest was grown in LB broth in a 28˚C shaker. The bacteria was then centrifuged for 10min at 4000RPM and resuspended in liquid MS medium 222 (Murashige and Skoog, 1962; Duchefa Biochemie ). The bacteria was then diluted to an OD of 0.4 at 660nm in liquid MS. Leaves from the plants grown in the magentas were cut to 1x1cm square sections and inserted to the agrobacterium, which was in 50ml falcon tubes. These Falcon tubes were then shaken for 15min at ~100RPM, after which leaf sections were removed and blotted on sterile Whatman filter paper and transferred to MS medium supplemented with 2mg/L 1-Naphthaleneacetic acid (NAA; Sigma Aldrich) and 0.2mg/L Kinetin (Sigma Aldrich) and placed in the dark for 48h. Leaf sections were then transferred to MS plates supplemented with 2mg/L Kinetin, 0.8mg/L Indole-3-acetic acid (IAA; Sigma Aldrich), 250mg/L Ticarcillin 2na & Clavulanate K (Duchefa Biachemie) and 100mg/ L Kanamycin monosulfate (Duchefa Biachemie). Kanamycin was used due to the plant selection being based on the *NPT2* Kanamycin resistance gene. This medium was replaced every two weeks until shoots developed on the leaf sections. Shoots were then cut and transferred to magentas containing MS medium supplemented with 250mg/L Ticarcillin and 100mg/L Kanamycin until root development, following which plants were transferred to hardening in soil.

*Whole plant drought trials*

Drought trials were performed during the summer of 2018, winter of 2019 and spring of 2020. plants were grown in cavity trays with potting soil in a growth room before being transferred for acclimation in the weighing lysimeter greenhouse. All parts of the system that would be placed on the scales were weighed, averaged and the average weight was fed to the system to calculate soil weight following pot filling. Pots were then filled with 4L potting soil that was saturated and homogenized by whirling in a cement mixer. A plaster cast was then inserted to keep soil capillary properties upon planting. Lysimeter scales were calibrated and pots with soil were placed on the system for initial measurement, after which plaster casts were removed and plants were transferred to the pots. Pots were then re-weighed to determine plant weight. These pots remained on the system for the duration of the experiment, with weight being recorded every 3 min. Plants were irrigated in excess in three pulses starting from 20:30 so irrigation derived alterations in pot weight would not interfere with daytime measurements. In the summer experiment, upon starting drought treatment plants were not irrigated during one day, following which they were irrigated with a volume of water equivalent to 50% of their previous day's transpiration as calculated by the system. Once each plant reached a daily transpiration rate below a daily transpiration threshold (which was set at approximately 20% of average daily transpiration the day prior to drought induction), irrigation was stopped for two additional days after which irrigation was fully resumed and recovery began. In the winter experiment, due to slow plant growth and low VPD, irrigation was stopped and only resumed when all plants were recovered simultaneously. The third (spring) experiment was performed with sand filled pots, which were not homogoniesed prior to pot filling, as this was un-necessary. In this experiment, plants were grown in the lysimeter greenhouse, 2.5 weeks prior to being placed on the system, in order to reach a significant transpiration rate prior to drought. Therefore, plant weight was determined by averaging the weight of 3-5 plants grown for this purpose alongside the plants which would ultimately be placed on the system. Drought was initiated with reduction to 50% of previous day's transpiration, and was stopped for two days after a plant transpired below the 20% transpiration threshold. Following these two days irrigation was resumed. In all three experiments, WUE was calculated as a plants biomass gain divided by its transpiration during the days prior to drought initiation. Plant dry biomass was weighed following at least three days of shoot drying in an oven heated to 60˚c. Theta points were calculated for each plant individually using a two piece linear curve, which takes average transpiration rates prior to drought, draws a non-sloped line at the average value of these points and then searches for the regression line of points representing declining transpiration under drought conditions which has the best $R^2$ values. Transpiration rate (E) values were calculated by dividing pot water loss by plant biomass as calculated by the system. Plant biomass was calculated by adding the difference between fully irrigated and drained pots' weight in consecutive nights.

**Mozo T, Hooykaas JJ** (1992) Factors affecting the rate of T-DNA transfer from *Agrobacterium tumefaciens* to *Nicotiana glauca* plant cells. Plant Mol Biol. 19: 1019-1030

**Murashige T, Skoog F** (1962) A revised medium for rapid growth and bioassays with tobacco tissue cultures Physiologia Plantarum **15**: 473–497
