## Supplemental figures S1-S14 for "Cuticular Wax Composition is Essential for Plant Recovery Following Drought with Little Effect under Optimal Conditions"

### Supplementary figures

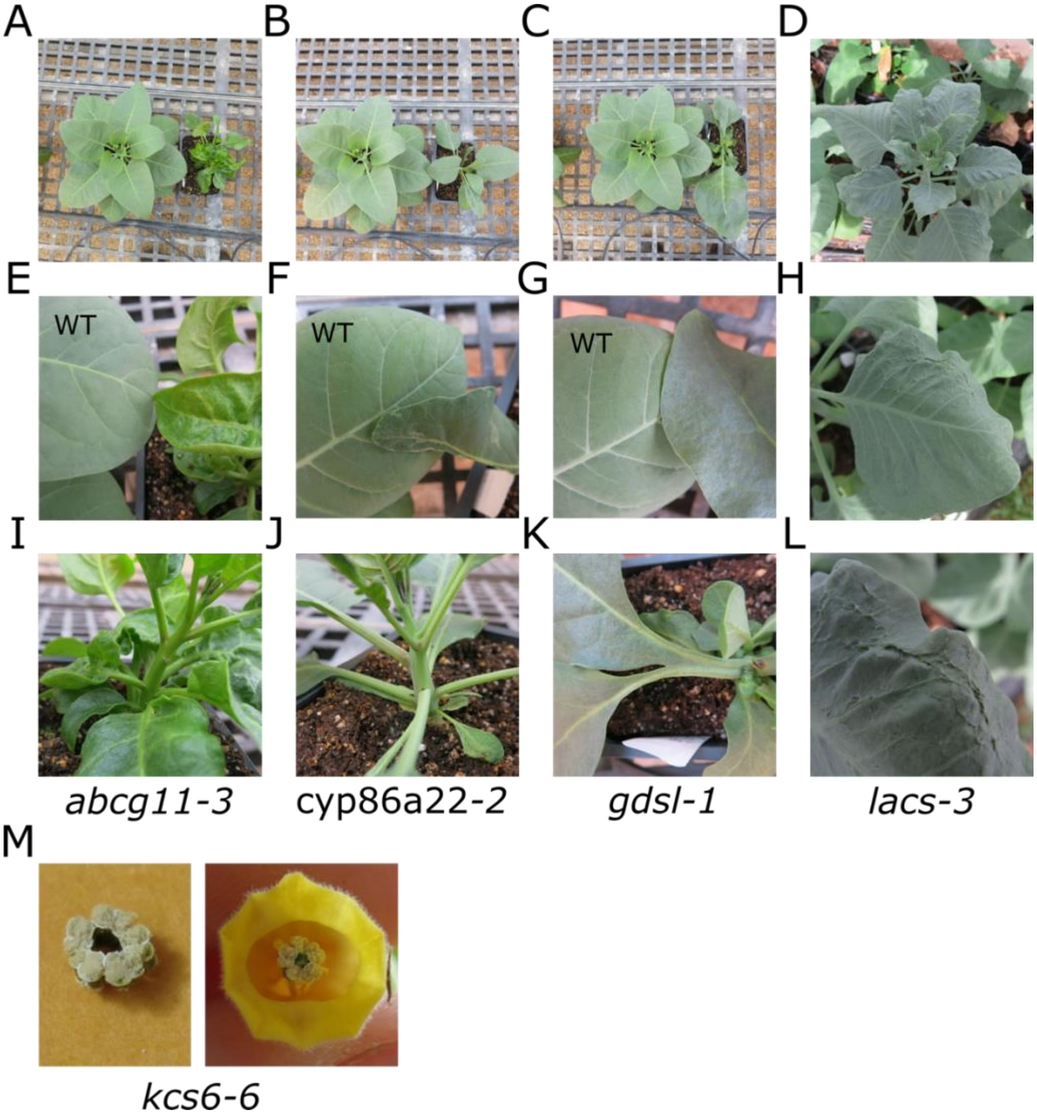


**Figure S1.** Images of additional CRISPR mutants displaying cuticular lipids metabolism- related phenotypes. Note glossiness and altered shapes of mutant leaves. **(A)** WT plant (left) beside an *abcg11-3* mutant. **(B)** A WT plant (left) beside a *cyp86-2* mutant. **(C)** A WT plant (left) beside a *gdsl-1* mutant. **(D)** A *lacs-3* mutant plant. **(E)** WT and *abcg11-3* leaves. **(F)** WT and *cyp86-2* leaves. **(G)** WT and *gdsl-1* leaves. **(H)** A *lacs-3* leaf. **(I)** An *abcg11-3* stem displaying a glossy appearance. **(J)** A *cyp86-2* stem displaying a glaucous appearance. **(K)** *gdsl-1* leaves displaying fused petioles. **(L)** A *lacs-3* leaf displaying wrinkles and cracking. **(M)** A fused anthers phenotype in *kcs6-6* flowers

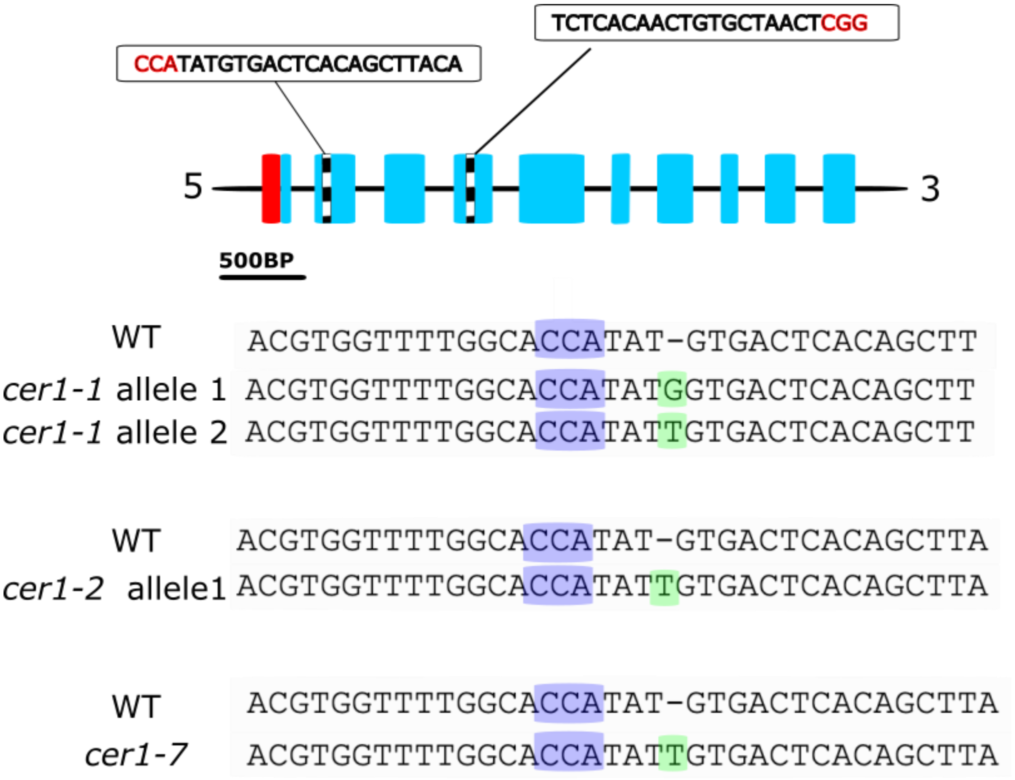


**Figure S2.** The *CER1* mutant alleles. The red boxes indicate UTRs, blue boxes represent exons, and black and white striped boxes indicate the locations of crRNA. The UTRs and exon sizes are correlated to their length; intron size is presented uniformly and is not correlated to the actual length. Purple shading on WT and mutant sequences indicates the PAM sequence. Green shading indicates an insertion.

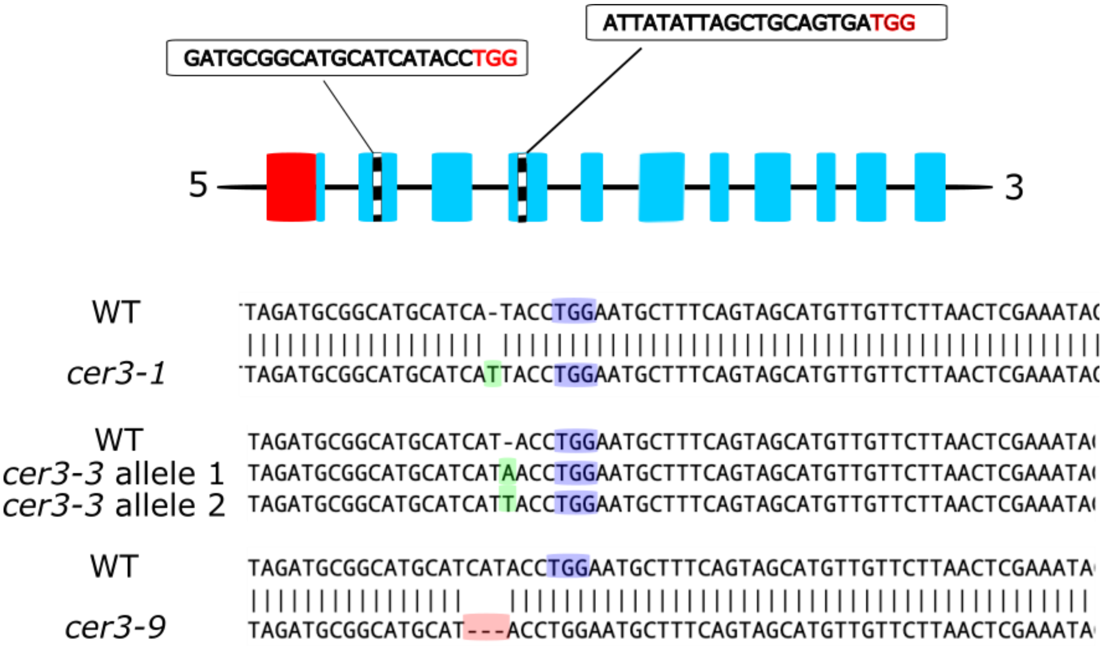


**Figure S3.** The *CER3* mutant alleles. The red boxes indicate UTRs, blue boxes represent exons, and black and white striped boxes indicate the locations of crRNA. The UTRs and exon sizes are correlated to their length; intron size is presented uniformly and is not correlated to the actual length. Purple shading on WT and mutant sequences indicates the PAM sequence. Red shading indicates a deletion and green an insertion.

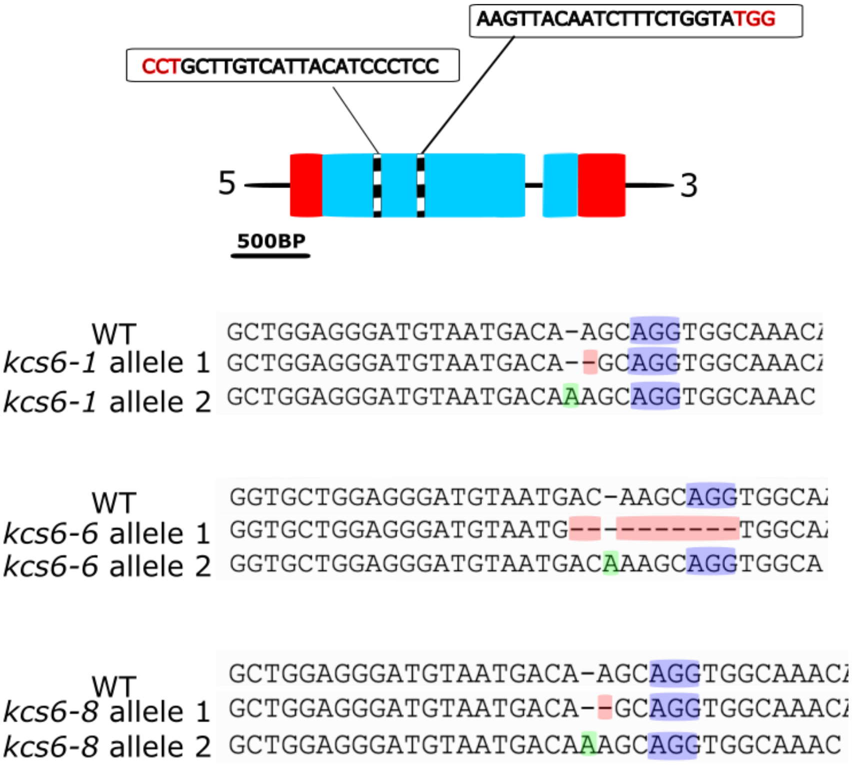


**Figure S4.** The *KCS6* mutant alleles. The red boxes indicate UTRs, blue boxes represent exons, and black and white striped boxes indicate the locations of crRNA. The UTRs and exon sizes are correlated to their length; intron size is presented uniformly and is not correlated to the actual length. Purple shading on WT and mutant sequences indicates the PAM sequence. Red shading indicates a deletion and green an insertion.

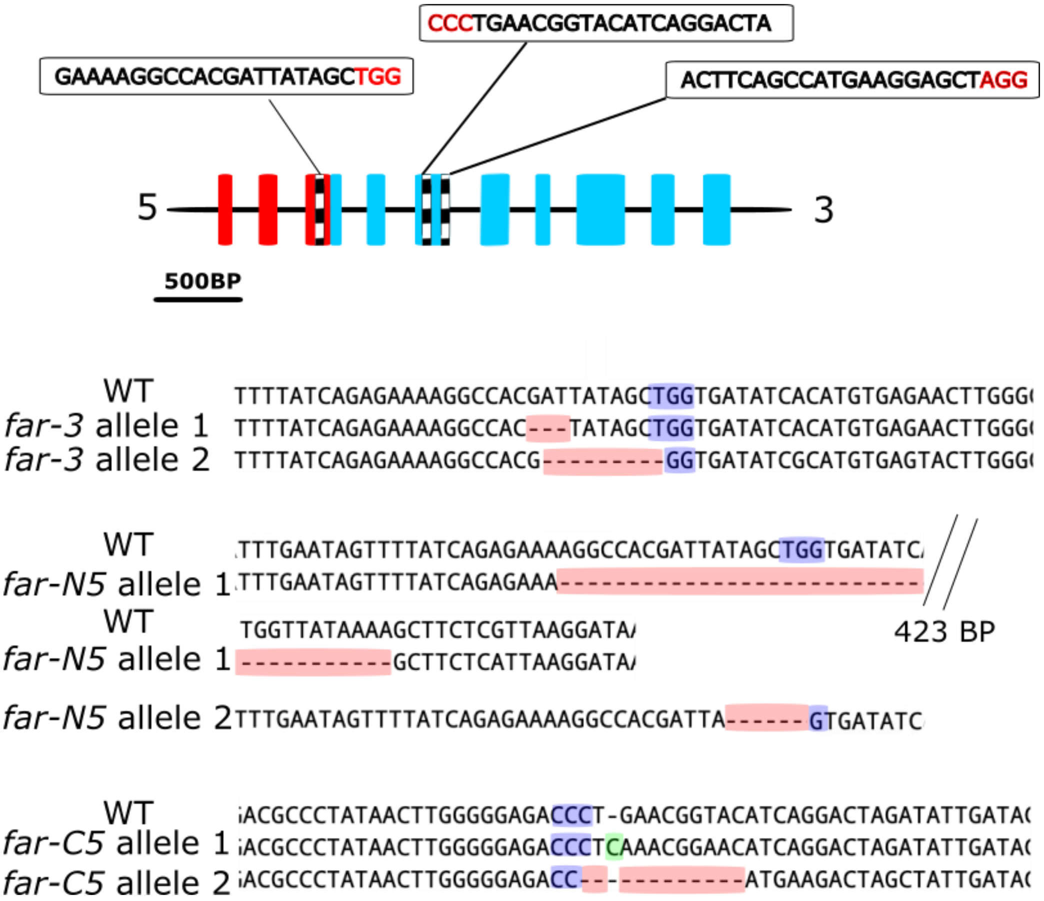


**Figure S5.** The *FAR* mutant alleles. The red boxes indicate UTRs, blue boxes represent exons, and black and white striped boxes indicate the locations of crRNA. The UTRs and exon sizes are correlated to their length; intron size is presented uniformly and is not correlated to the actual length. Purple shading on WT and mutant sequences indicates the PAM sequence. Red shading indicates a deletion and green an insertion.

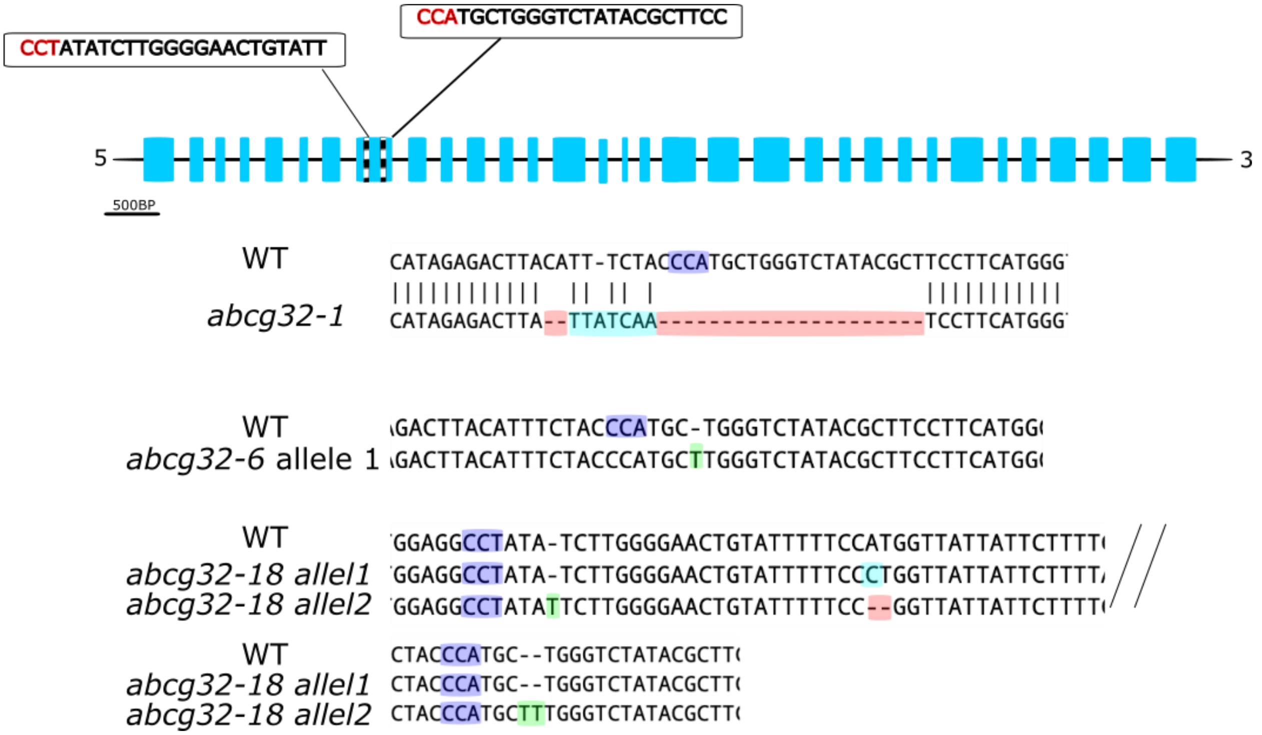


**Figure S6.** *ABCG32* mutant alleles. Blue boxes represent exons, and black and white striped boxes indicate the locations of crRNA. The exon sizes are correlated to their length; intron size is presented uniformly and is not correlated to the actual length. Purple shading on WT and mutant sequences indicates the PAM sequence. Red shading indicates a deletion, green shading indicates an insertion and blue a substitution.

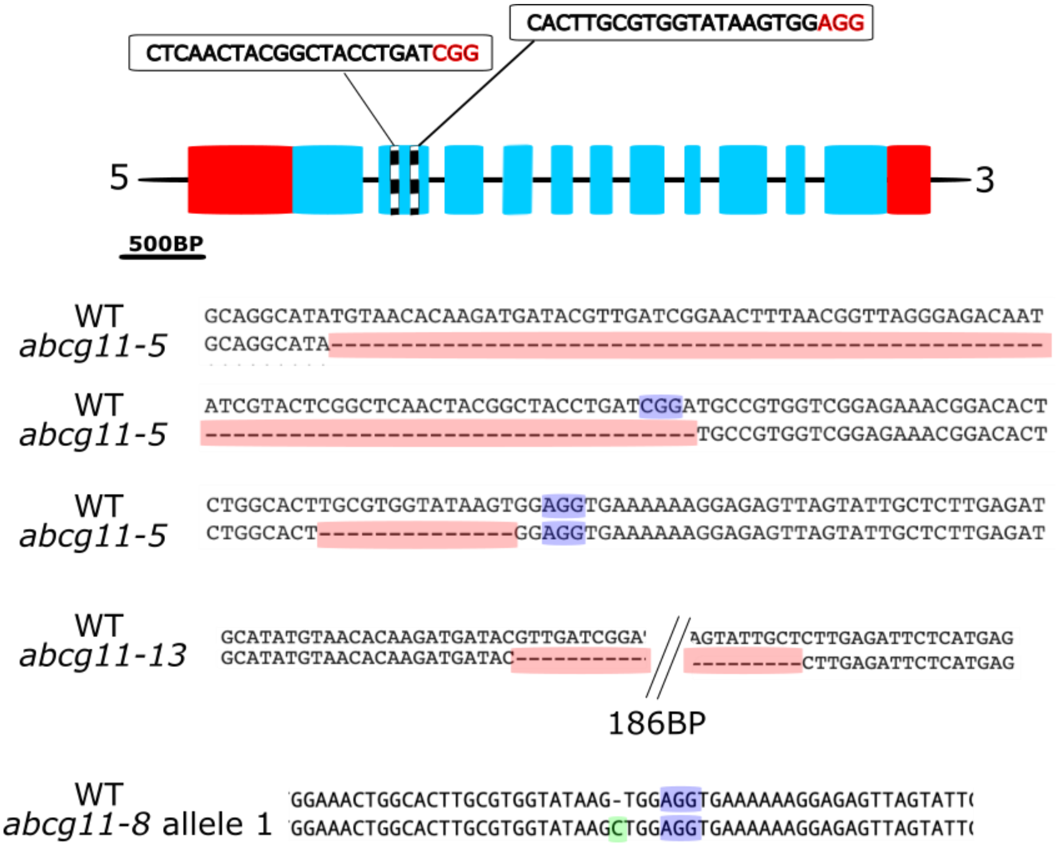


**Figure S7.** *ABCG11* mutant alleles. The red boxes indicate UTRs, blue boxes represent exons, and black and white striped boxes indicate the locations of crRNA. The UTRs and exon sizes are correlated to their length; intron size is presented uniformly and is not correlated to the actual length. Purple shading on WT and mutant sequences indicates the PAM sequence. Red shading indicates a deletion and green shading an insertion.

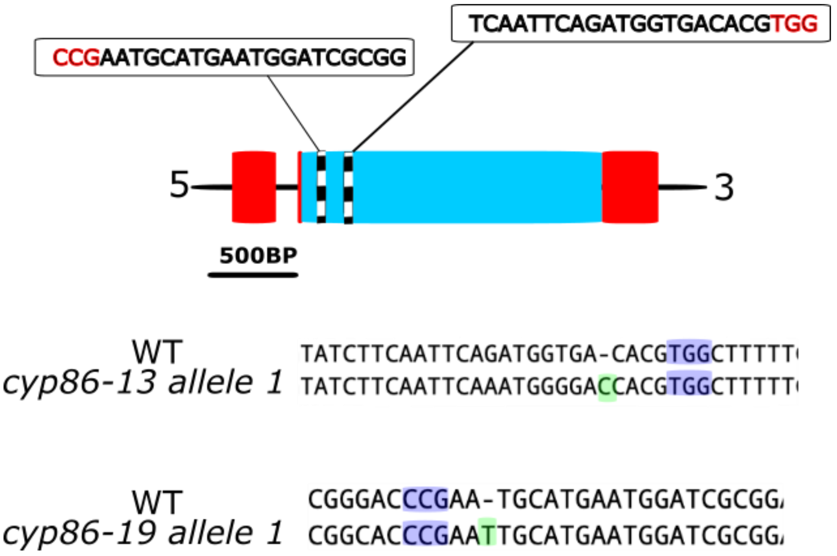

**Figure S8.** *CYP86A22* mutant alleles. The red boxes indicate UTRs, blue boxes represent exons, and black and white striped boxes indicate the locations of crRNA. The UTRs and exon sizes are correlated to their length; intron size is presented uniformly and is not correlated to the actual length. Purple shading on WT and mutant sequences indicates the PAM sequence.. Green shading indicates an insertion.

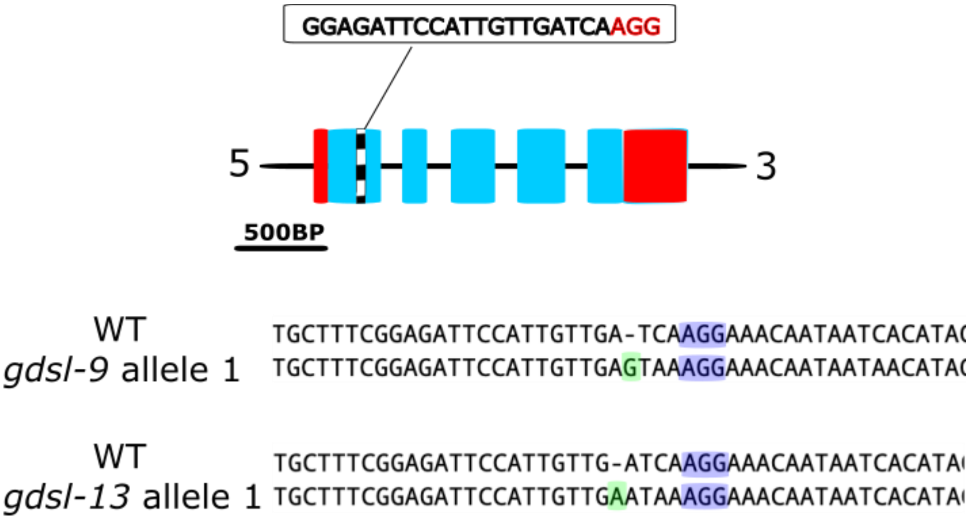


**Figure S9.** *GDSL* mutant alleles. The red boxes indicate UTRs, blue boxes represent exons, and black and white striped boxes indicate the locations of crRNA. The UTRs and exon sizes are correlated to their length; intron size is presented uniformly and is not correlated to the actual length. Purple shading on WT and mutant sequences indicates the PAM sequence. Green shading indicates an insertion.

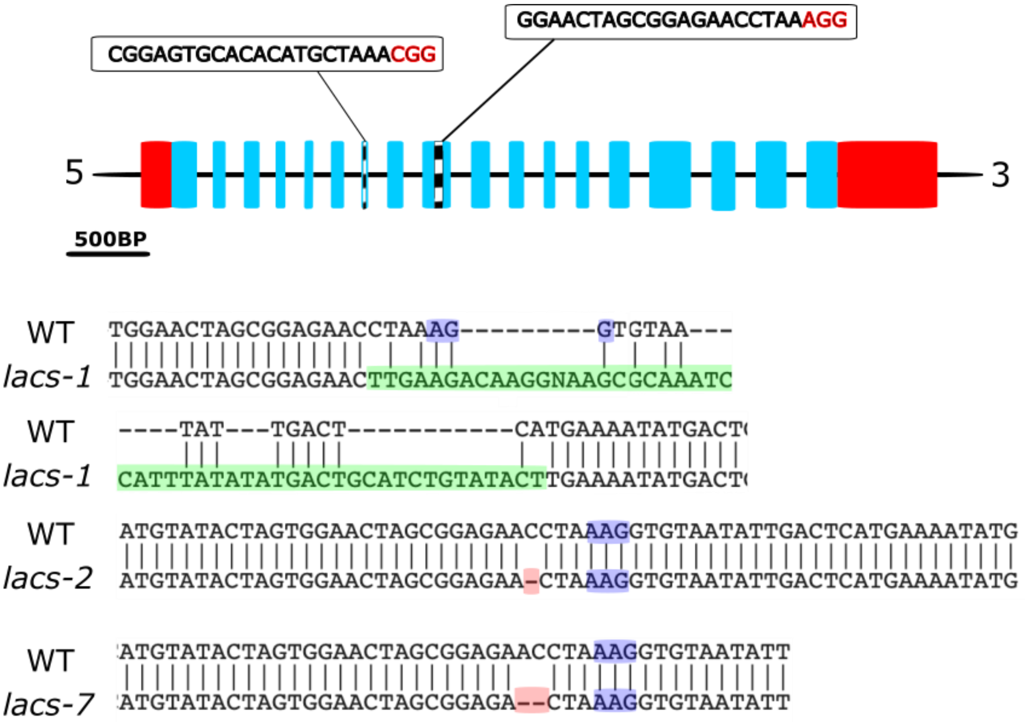


**Figure S10.** *LACS* mutant alleles. The red boxes indicate UTRs, blue boxes represent exons, and black and white striped boxes indicate the locations of crRNA. The UTRs and exon sizes are correlated to their length; intron size is presented uniformly and is not correlated to the actual length. Purple shading on WT and mutant sequences indicates the PAM sequence. Red shading indicates a deletion, green shading indicates an insertion.

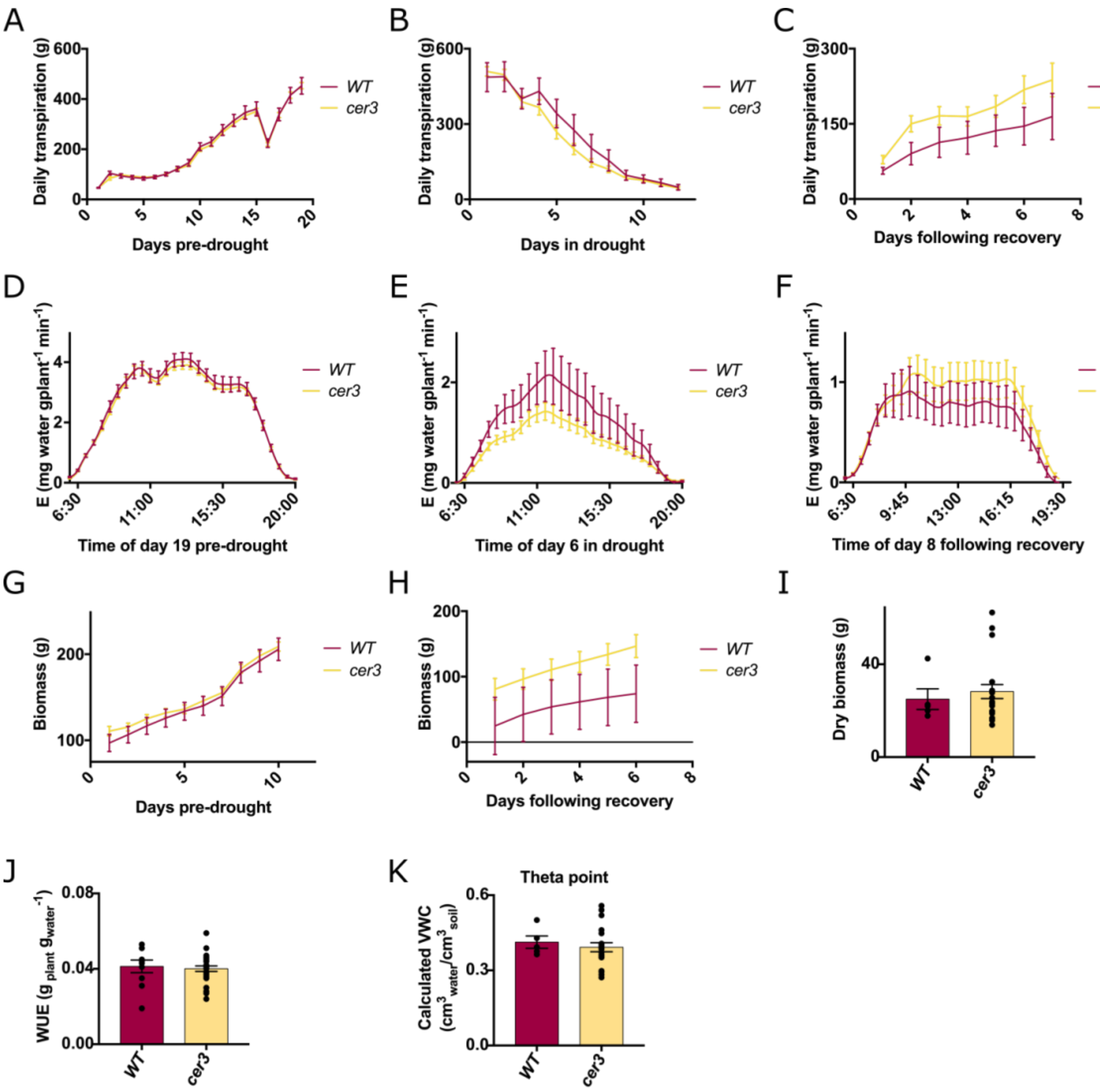


**Figure S11.** Multiple parameters assays of *cer3* plants response. Plants grown during the summer of 2018. **(A)** Pre-drought daily transpiration. **(B)** Daily transpiration following drought initiation. **(C)** Daily transpiration following recovery. Since each individual plant received full irrigation three days after it transpired below the 20% daily transpiration threshold (see materials and methods), the days in recovery unlike those pre-drought and during drought are different for each individual plant. **(D)** Transpiration rate during day 19 prior to drought induction. Measurements were taken every 3 minutes, though error bars are only shown every 30 minutes. **(E)** Transpiration rate during the sixth day of drought. **(F)** Transpiration rate during the day which in average was eight days following recovery (the exact time post recovery varied from plant to plant). **(G)** Plant biomass during the period prior to drought induction. **(H)** Plant biomass during the six days before the experiment's end, at which time all plants had been resuscitated. **(I)** Dry shoot biomass at the experiments end. **(J)** WUE as calculated pre-drought. **(K)** "Theta point"; volumetric water content at which plants began reducing transpiration rate in response to drying soil. In WT n= 5 to 10 and *cer3* n= 20 to 29 plants. All *cer3* data is the average of lines derived from two independent mutant alleles (i.e. *cer3-1* and *cer3-9*). All bars represent standard errors.

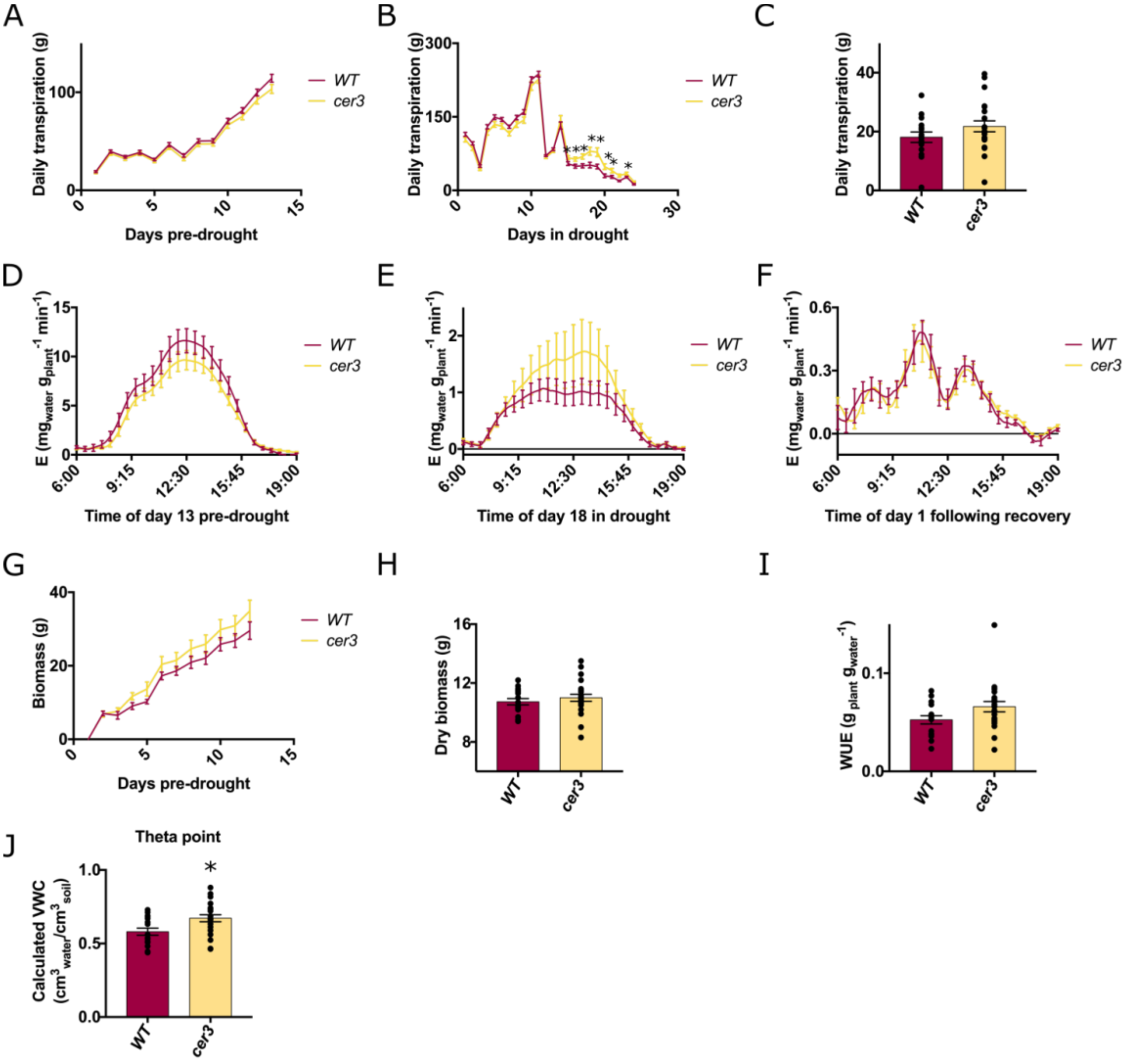


**Figure S12.** Multiple parameters assays of *cer3* plants response. Plants grown during the winter of 2019. **(A)** Pre-drought daily transpiration. **(B)** daily transpiration following drought initiation. **(C)** Daily transpiration following recovery. All plants were recovered simultaneously due to low transpiration rates, which led to many plants not reaching below the 20% daily transpiration threshold. **(D)** Transpiration rate during day 13 prior to drought induction. Measurements were taken every 3 minutes, though error bars are only shown every 30 minutes. **(E)** Transpiration rate during the 18th day of drought. **(F)** Transpiration rate during the day of recovery. **(G)** Plant biomass during the period prior to drought induction. **(H)** Dry shoot biomass at the experiments end. **(I)** WUE as calculated pre-drought. **(J)** "Theta point"; volumetric water content at which plants began reducing transpiration rate in response to drying soil. In WT n= 14 to 16 and *cer3* n= 20 to 23 plants. All *cer3* data is the average of three independent mutant alleles (*cer3-1*, c*er3-3* and *cer3-9*). All bars represent standard errors. Asterisks indicate significance of $p<0.05$ as determined by a student's t test.

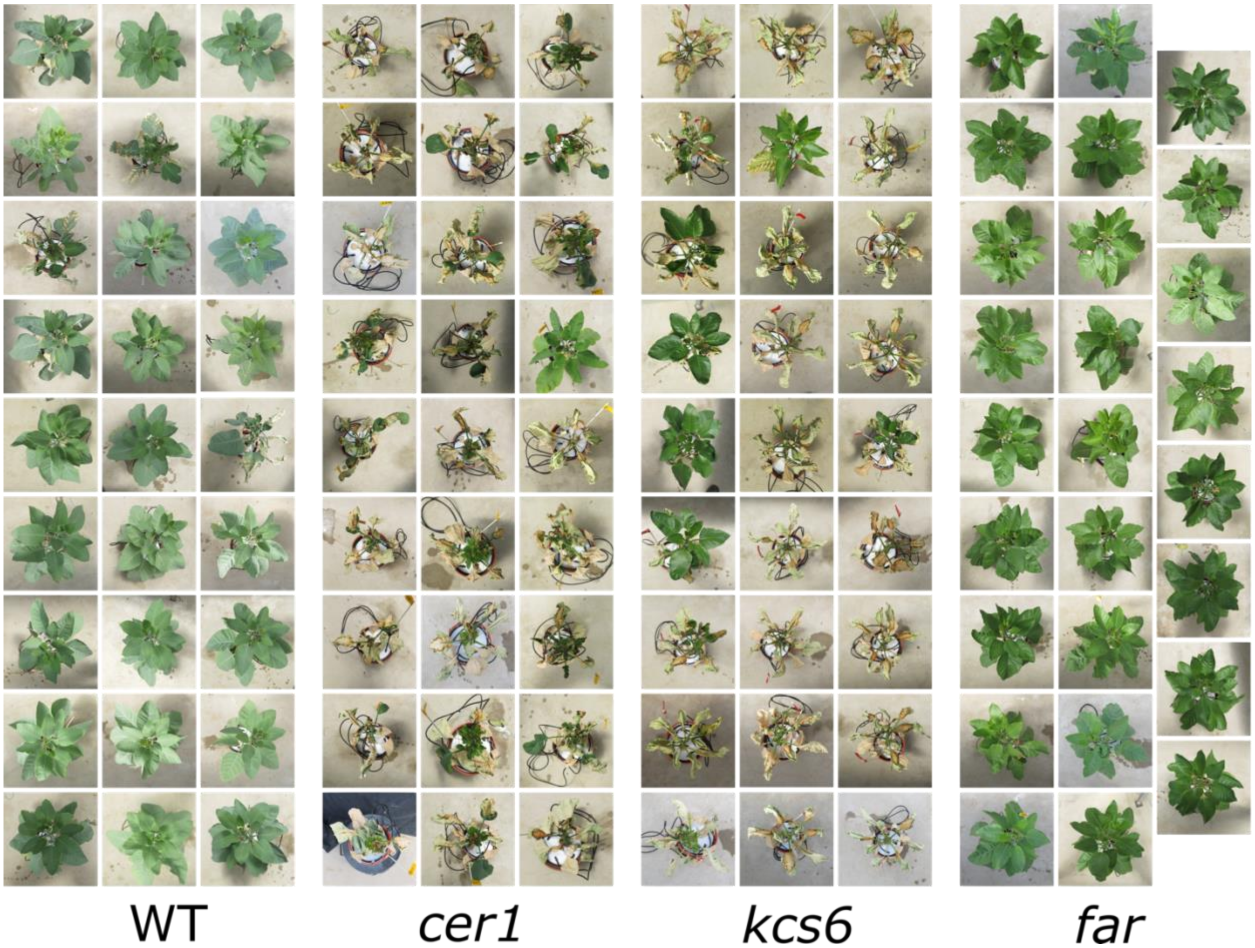


**Figure S13.** Photographs of plants at the end of the spring drought experiment. Plants are clustered by the gene they are mutated in. It may be seen that while *far* mutants all maintained a green, non-dried phenotype, a great majority of *cer1* and *kcs6* mutants had leaves dry out during the drought treatment. WT plants show a majority of plants with green non-dried leaves though there are several plants which experienced leaf death during drought.

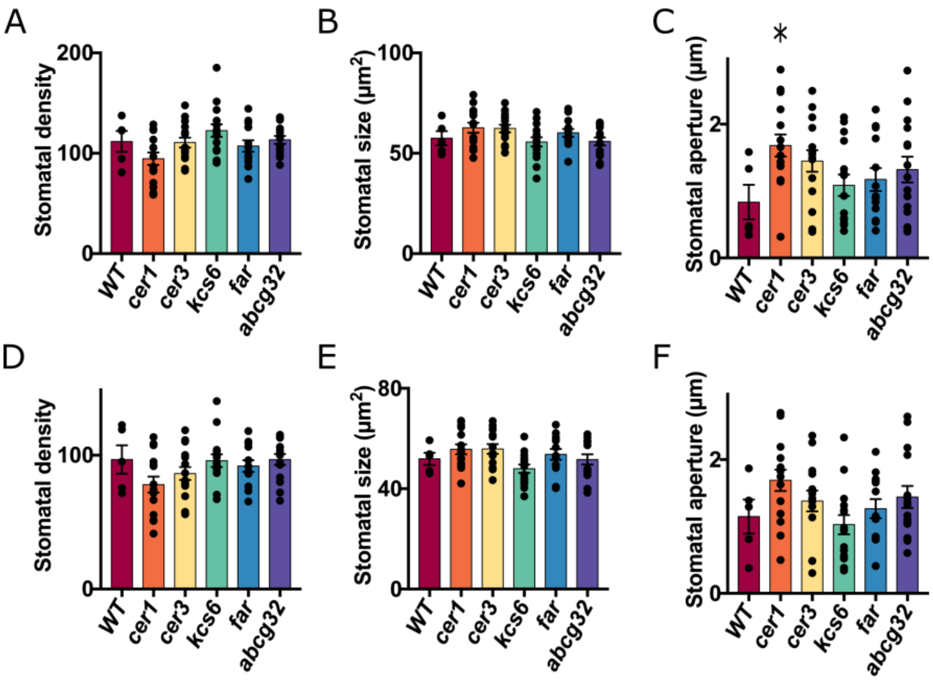


**Figure S14.** Effect of epicuticular wax composition in the different cuticular wax mutants on stomatal density size and aperture. **(A-C)** Abaxial side stomata. **(D-F)** Adaxial side stomata. **(A, D)** Stomatal density per 0.1mm$^2$; **(B, E)** stomatal size. **(C, F)** stomatal aperture. WT n=5, other lines n=15. Different genes data is composed of three independent lines mutated in that gene. Each biological replicate included photography and analysis of two fields for each imprint. For stomatal size and aperture between 3-10 stomata were analyzed per field, depending on stomatal density and quality of focus. These measurements were then averaged. Averages of two fields in each imprint were averaged as well. Bars represent standard errors. Asterisks indicate a significance of $p<0.05$ as determined by a student's t test.
